# Principles of Local and Global Grouping that Underlie Segmentation of Natural Texture Images

**DOI:** 10.64898/2026.05.06.723304

**Authors:** Wilson S. Geisler, Abhranil Das

## Abstract

The human visual system segments images using both high-level recognition mechanisms and low-level mechanisms that are largely independent of specific prior experience. The low-level mechanisms are essential for initiating recognition processes, and for learning to recognize new materials, objects, and contexts. Here we describe a hierarchical Bayesian observer (HBO) model of texture segmentation that is biologically plausible, takes into account the statistics of natural scenes, and does not depend on prior experience. The HBO model consists of five steps: local similarity grouping with local normalization, mutual similarity grouping (local grouping is strengthened if the neighboring regions are similar to the same set of other regions), transitive grouping (good continuation), confidence grouping (neighboring regions far from the same-different decision boundary guide grouping of regions near the decision boundary), and region grouping (similarity grouping of the regions from the initial segmentation). We find that a local similarity grouping process, trained to maximize accuracy, predicts human texture discrimination accuracy. We then find that the four additional steps accurately segment images with randomly shaped regions containing arbitrary natural textures. The success of the model depends on all the steps, but especially on local-similarity and transitive grouping. We also find that the transitive grouping allows correct segmentation of non-stationary texture regions (e.g., textures slanted in depth). Further, we find that when illumination varies across the image, local normalization enables both correct texture segmentation and estimation of illumination change. Finally, we find that unlike our model large state-of-the-art deep networks often fail on these stimuli.

## Introduction

Humans (and many other animals) have a remarkable ability to segment natural scenes into physically meaningful regions. The segmented regions often correspond to different materials, different kinds of illumination, different objects, or different connected parts of objects. The representation of these regions is often hierarchical. For example, a segmented region corresponding to a specific object may simultaneously be segmented into regions corresponding to different illuminations and different object parts.

The segmentation process is very sophisticated and contains many components. While some of these components are “high level”, exploiting recognition of materials, objects and/or scene context, others are more “low level” and largely independent of specific prior experience. The upper images in Fig. 1 show examples of images containing five randomly shaped regions each filled with a different randomly selected natural texture. The human visual system has little trouble correctly segmenting such images even though the region shapes (lower images in Fig. 1) and the textures are largely unfamiliar. In this paper, the focus is on such low-level mechanisms that allow accurate segmentation of images like those in Fig. 1.

**Figure 1.**
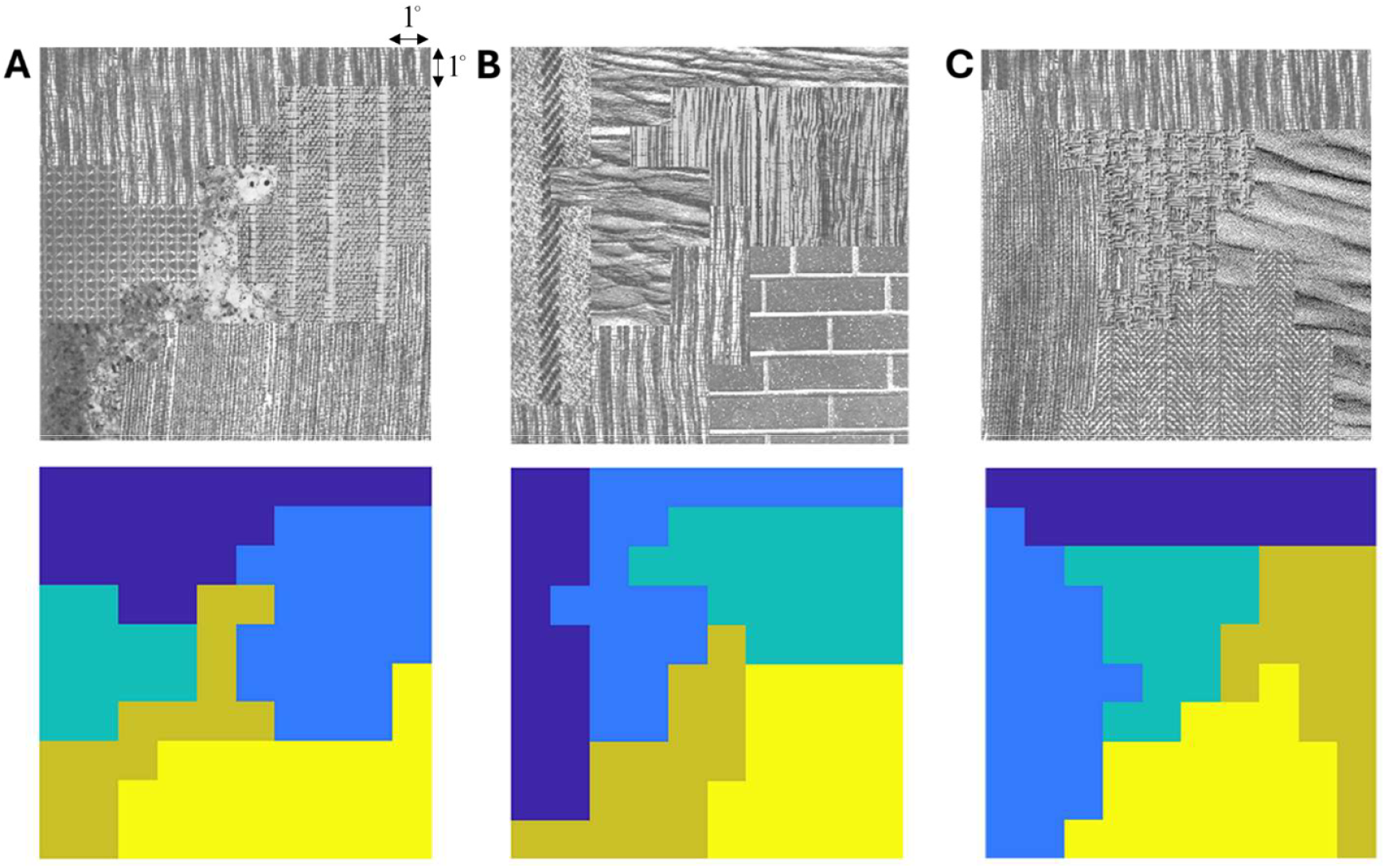
Grown texture region (GTR) images (see later section for a description of how these images are created). Upper row: random texture regions filled with samples from Brodatz textures. Lower row: the ground-truth texture regions are indicated with a uniform color. These are also the regions found by the hierarchical Bayesian observer (HBO) segmentation model. Number of regions = 5, block/patch size = 1°.

The low-level mechanisms are absolutely essential, in that they are required for initiating recognition processes and for making possible the learning of new objects and materials. In natural scenes the low-level mechanisms usually provide a segmentation that is sufficient to trigger recognition of objects, materials, and context, which can then provide feedback or recurrent signals to produce a more accurate segmentation. We will argue later that the low-level components of segmentation also play an important role in recovering variations in illumination as well as the 3D geometry of surfaces from perspective cues.

The perceptual consequences of many of the low-level components were first highlighted by the Gestalt psychologists’ demonstrations of perceptual grouping (Wertheimer 1938; 1961). Many of the demonstrations show the importance of similarity. All other things being equal, local clusters of features that are more similar in color and geometry are more likely to be grouped together and segmented from less similar clusters of features. Other demonstrations show the importance of continuity, symmetry, and parallelism. Feature properties that change smoothly over space tend to be grouped into larger clusters. The classic example is contour grouping by good continuation (e.g., Elder & Goldberg 2002; Field et al. 1993; Geisler et al. 2001). Similarly, contours that are symmetric or parallel along some axis tend to be grouped together (Wagemans 1997). These Gestalt principles of grouping have been studied extensively, but usually separately, and with relatively simple discrete-element stimuli (for reviews see Wagemans et al. 2012a, Wagemans et al. 2012b, Palmer 1999).

The Gestalt principles are sensible in that they correspond to physically meaningful properties of natural scenes. Similar clusters of features that are nearby tend to be part of the same object or to be part of the same surface. Biological growth tends to produce continuous boundary contours that are locally approximately symmetric or parallel. Even erosion together with gravity tends to produce physically symmetric or parallel structures.

The Gestalt principles have been incorporated into computational and computational-neuroscience studies of texture and scene segmentation. Early computational models incorporated biologically plausible mechanisms such as spatial-frequency and orientation tuned filters followed by simple nonlinearities and additional filtering (for review see Landy & Graham 2004). These models use similarity and discontinuities in spatial-frequency and orientation content to identify different texture regions. The models are able to account for a number of aspects of human texture segmentation performance for simple texture stimuli. However, they were not designed to take into account the statistical properties of natural texture images.

More recent computational models have shifted toward using large-scale machine learning and have been mostly directed toward the general problem of natural scene segmentation, and toward practical application, and hence have not been constrained to biologically plausible computations (Minaee et al. 2022; Kirillov et al. 2023). Because they are trained on natural images, they incorporate some natural image statistics. However, they depend strongly on recognition of context and scene content and are not designed to segment images like those in Fig. 1, which do not contain recognizable context or content. We show later that these models often do not do as well on these stimuli.

The approach taken here is to work toward the development of image computable hierarchical Bayesian observer (HBO) models that are biologically plausible, exploit the statistical properties of natural scenes, and do not depend on prior experience (recognition). An overview of the proposed framework is shown in Fig. 2 (details are given in the Results and Appendix). The first step is local similarity grouping of neighboring small image regions across the visual field. This grouping rule combines measurements of natural image statistics and Bayesian likelihood computations with optimal decision variables trained on natural texture. The second step is mutual similarity grouping which strengthens local grouping if the two neighboring regions are similar to the same set of other regions in the image. The third step is transitive grouping which implements the Gestalt principle of continuity (good continuation) on the local regions: if region ***a*** groups with neighboring region ***b***, and region ***b*** groups with neighboring region ***c***, then regions ***a*** and ***c*** are grouped. This step provides an initial segmentation of the image. The fourth step is confidence grouping. During step 2 local similarities that are near the same-different decision boundary are identified and left unlinked. After step 3 they may be assigned to a region based on their similarity to the local regions within the initial groups. The fifth step is region similarity grouping. The similarity of the groups from step 4 are used to decide which, if any, groups should be merged. The segmentations shown in the bottom row of Fig. 1 were generated from the images in the upper row of Fig. 1 by implementing the steps in Fig. 2.

**Figure 2.**
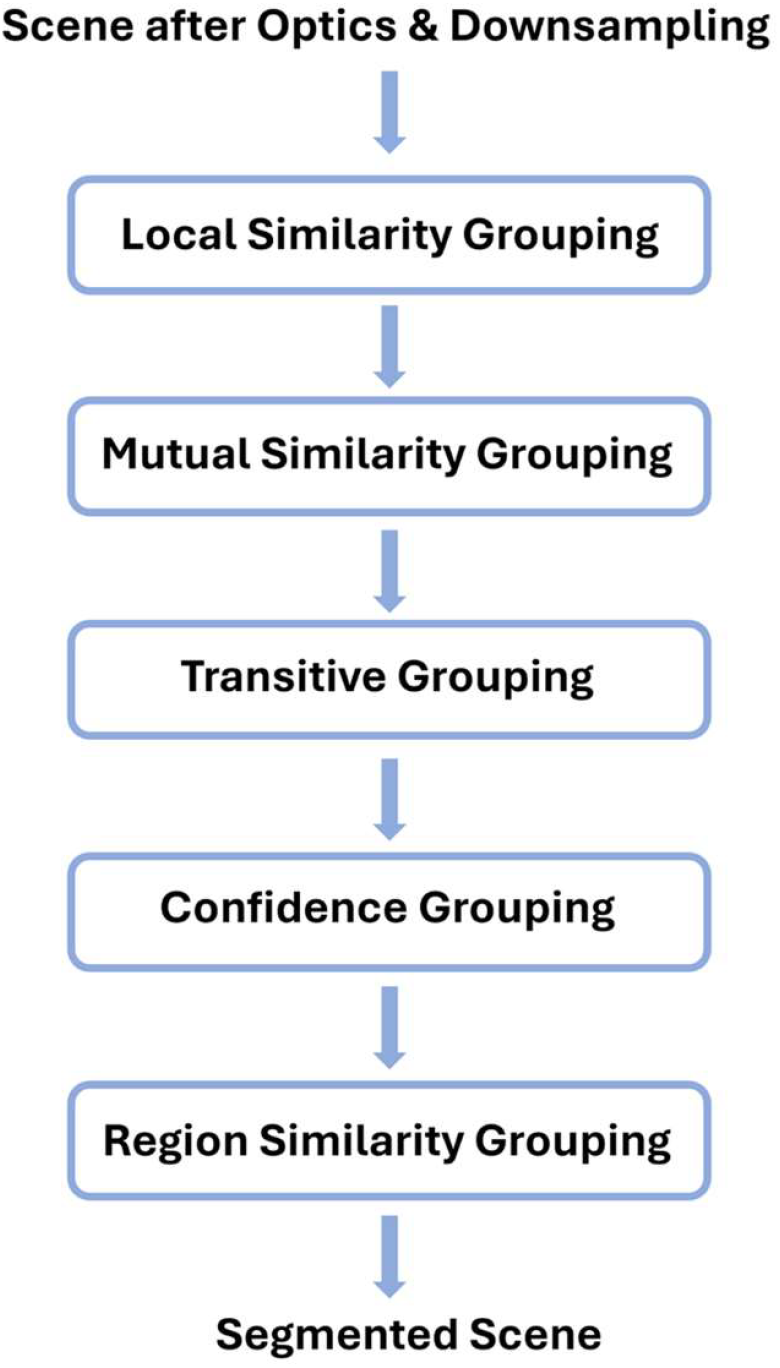
Hierarchical Bayesian observer (HBO) framework of image segmentation models.

The first section below describes the theory, natural-scene statistics, and human experiments underlying an HBO model of texture discrimination. The decision variable for texture discrimination is the input to local similarity grouping, which is the foundation of the whole approach. The second section describes the rest of the steps in the HBO model of texture segmentation as well as the performance of the full model.

## Results: Texture Discrimination

The purpose of local similarity grouping is to determine the likelihood that neighboring image patches are from the same category or different categories of texture, and then to link the two patches together if the likelihood that they are from the same category is high enough. We focus here on 64×64-pixel patches that are displayed at a visual angle of approximately 1 deg. Thus, the range of spatial frequencies contained in the patches is from approximately 1 c/deg to 32 c/deg. We picked this scale for computational simplicity and because 1 c/deg is roughly the peak spatial frequency of the largest receptive fields in early visual cortex that encode the fovea and near periphery. The images in Fig. 1 are composed of 100 patches. In other words, we are considering displays where the texture images subtend 10 × 10 deg of visual angle. The approach we are taking can be generalized to other image sizes and patch sizes.

To develop and test principled models of local similarity grouping we consider the simple same-different discrimination tasks illustrated in Fig. 3. On each trial the observer fixates a small square, and then a pair of patches are briefly presented. The observer reports whether the two patches are samples from the same large sheet of texture or are from different sheets. In the “joined” task (Fig. 3A), the patches are abutting. In this task, when the patches are “same,” they are a single randomly sampled patch that is twice as long as wide. When they are “different” they are randomly sampled from different sheets (as shown here). In the “separated” task (Fig. 3B), the patches are separated from each other. In this task, when the patches are “same”, they are randomly sampled from the same sheet (as shown here) and when they are “different” they are randomly sampled from different sheets. The task in Fig. 3A is most relevant to local similarity grouping, whereas the task in Fig. 3B is most relevant to some of the later steps in Fig. 2. We develop a model for these two tasks and then compare its performance to that of human observers in the same tasks. The parameter-free predictions of the model were derived before measuring human performance.

**Figure 3.**
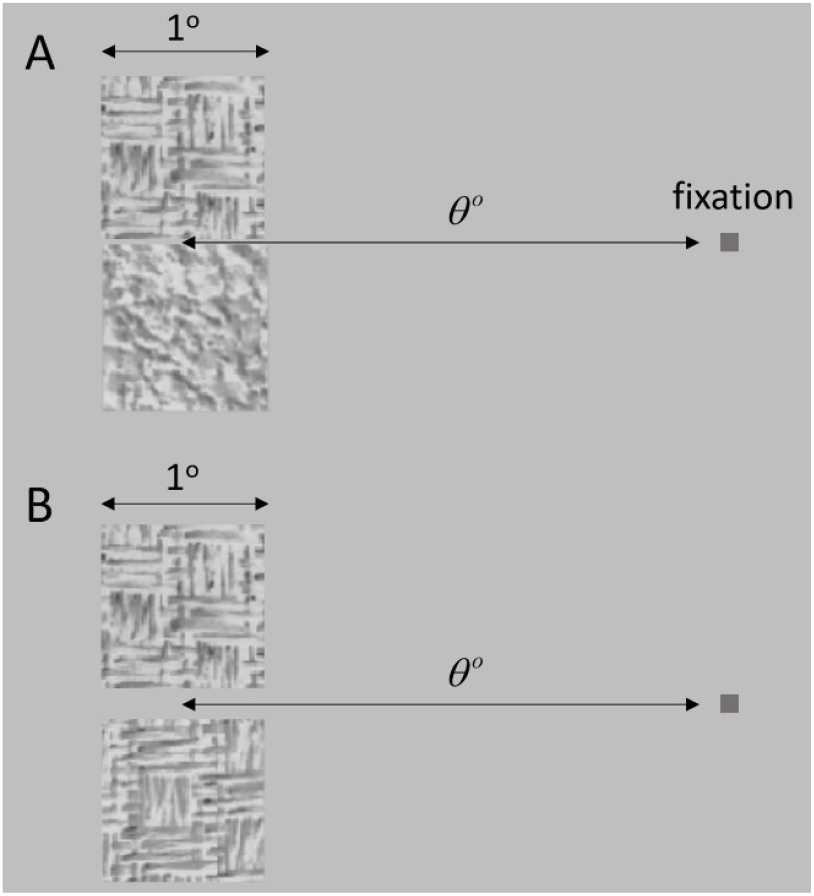
Same-different texture discrimination tasks. **A**. Joined task. **B**. Separated task.

A popular texture discrimination task is the “odd-ball” task (e.g., Freeman et al. 2013; Lieber et al. 2023) where three texture patches are presented, and the observer reports which pair is most similar. This is a simpler task because the observer does not have to make an absolute judgment of whether a pair of patches are samples from the same texture sheet. However, texture segmentation under natural conditions requires making absolute judgments of whether or not to bind local regions together, and thus the simple same-different task is more appropriate for our aims.

### Hierarchical Bayesian Observer Model for Texture Discrimination

Fig. 4 outlines a principled theoretical framework for the two discrimination tasks in Fig. 3, as well as for many other perceptual tasks (the steps for segmentation in Fig. 2 will be described later). The first step (Fig. 4A) is to specify a set of image features to extract, then determine their joint distribution in natural images, and then, if appropriate, transform the axes so the joint distributions have small correlations. The aim of this first step is to measure the prior distribution of the feature responses largely independent of the specific task. The idea is that evolution and learning over the lifespan will exploit the prior distribution when evolving or learning task-specific features. The task-independent priors do not solve the task, but they are easy to measure, and they provide useful constraints for defining and implementing the task-specific features. The second step (Fig. 4B) specifies the task-specific features (computed from the input feature responses), then uses the image-feature prior and principles of optimal encoding when deriving the computations that give the task-specific feature likelihoods and their joint log likelihood distribution. The third step (Fig. 4C) combines the log-likelihood distribution with a cost function and task-specific prior in order to determine the general decision rule. This defines the HBO model. The fourth step (Fig. 4D) is to simulate the performance of the model.

**Figure 4.**
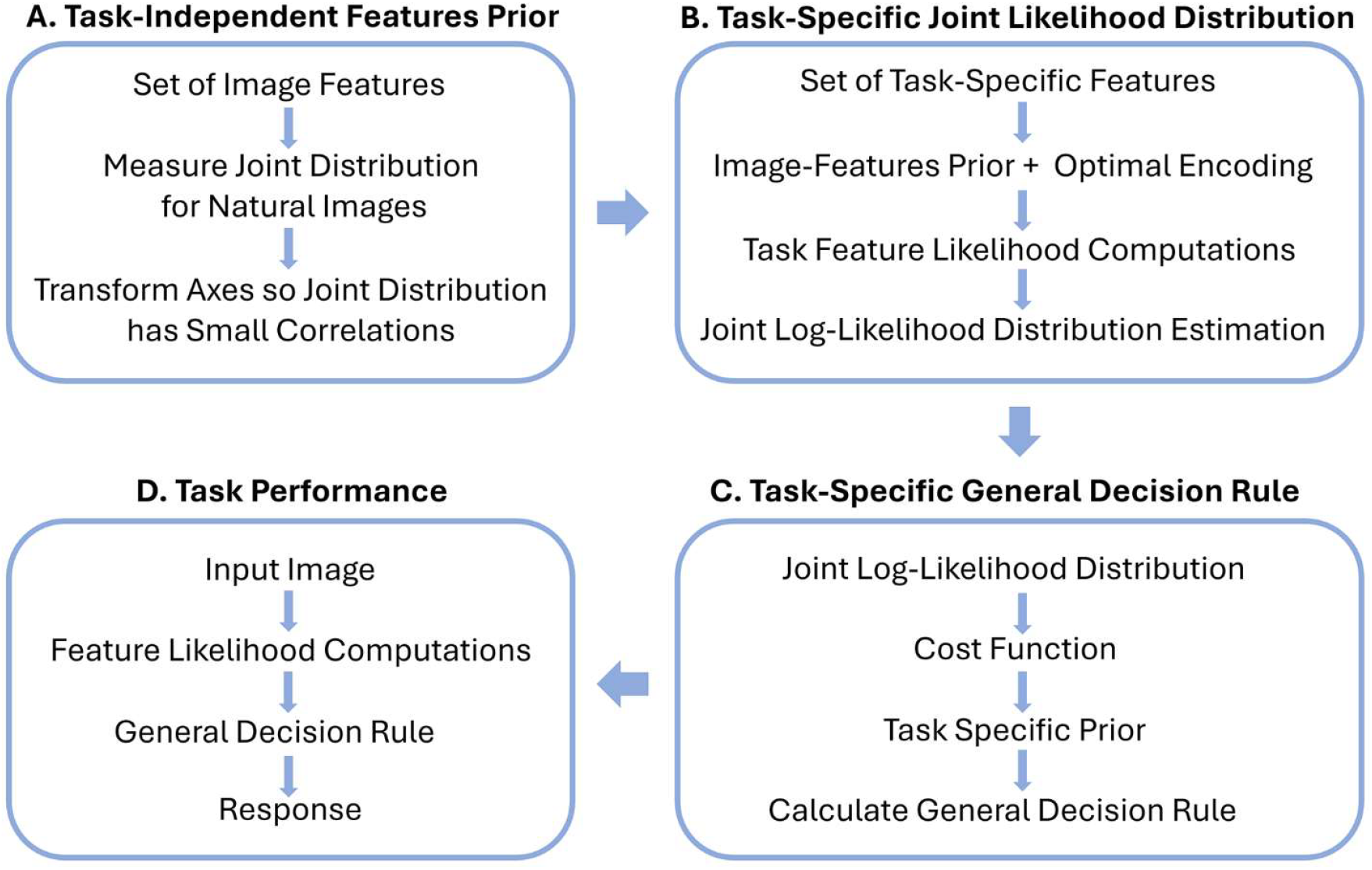
Hierarchical Bayesian Observer (HBO) framework for local texture discrimination.

We now lay out some of the details of the HBO model for the same-different texture discrimination tasks. The proposed input features are traditional and biologically plausible: (1) the set of long, middle, and short wavelength (LMS) cone responses to each pixel, (2) the set of first and second order steerable spatial filter responses, which captures the information in the cortical simple cell responses, (3) non-orientation-selective center-surround filters at two scales that also captures information from another subset of cortical simple cells, and (4) the power spectrum of the 1×1 deg patch, which captures the information in the cortical complex cell responses. Importantly, each 1×1 deg patch is mean and contrast normalized. The normalization values for each patch are saved for use in other computations (see Discussion), although they are not used in the current task. For details see the Appendix.

#### Task independent priors

To measure the task-independent features prior for the cone and spatial-filter responses, we measured the joint distribution of feature responses for patches randomly sampled from 391 natural images, captured with a calibrated high resolution (4282 × 2844), 14-bits per color, digital camera (see https://natural-scenes.cps.utexas.edu/). The images include outdoor scenes with no humans or human-made objects, as well as indoor and outdoor scenes with human-made objects. Before measuring feature responses, we applied the average human optical transfer function (OTF) to each natural image. We then converted each image from the camera’s RGB color space to the human LMS color space based on our measurements of the spectral sensitivities of the camera’s sensors.

The joint distributions were measured separately for the first two kinds of feature responses. For the cone responses, we first measured the joint distribution of the L, M and S values and then performed PCA to estimate the approximately orthogonal axes (which are close to being the classic achromatic, blue-yellow, and red-green color axes). Fig. 5A shows the marginal distributions of LMS values for the natural images. Fig. 5B shows the marginal distributions of the achromatic, blue-yellow, and red-green values. The joint distribution is approximately the product of these three marginals. Fig. 5C shows the cumulative probability distribution for the three distributions in Fig. 5B. These prior distributions are useful for optimizing the task-specific features.

**Figure 5.**
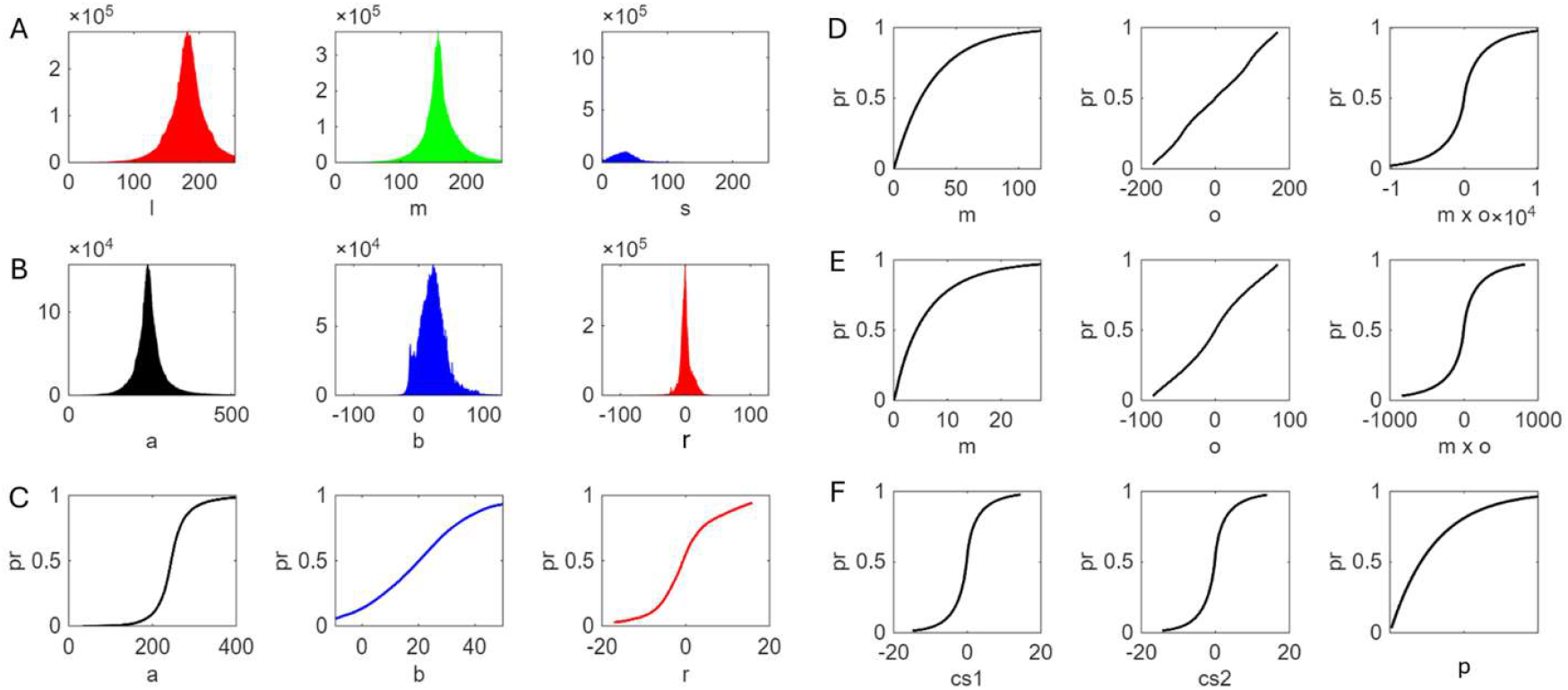
Task independent priors for color and contour features. **A**. Marginal histograms of *lms* cone responses. **B**. Marginal histograms of the PCA component *abr* (opponent channel) responses to all the natural images. **C**. Cumulative distribution functions (CDFs) for opponent color dimensions. **D**. CDFs for magnitude and orientation of 1^st^ derivative Gaussian steerable filter responses. **E**. CDFs for 2^nd^ derivative Gaussian steerable filter responses. **F**. CDFs for the small and large center surround filter responses, and for the power at any given spatial frequency in natural image patches (p axis range varies with the patch and spatial frequency).

For the orientation-selective spatial-filter (simple-cell) responses we computed, for each pixel location in the randomly sampled patches, the filter orientation with the largest response. We did this separately for the 1^st_^ and 2^nd^-derivative steerable filters. The steerable filters were as small as possible: 3 × 3 pixels. Even at this scale they accurately measured responses for all orientations (see Appendix). We found that small filters capture the most texture information from the small 1° x 1° patches of texture. Figs. 5D and 5E show the cumulative marginal probability distributions for response magnitude, orientation, and the product of magnitude and orientation. The product distribution describes the covariance between the magnitude and orientation responses. Fig. 5F shows the cumulative distributions for the small (3 × 3) and large (5 × 5) center-surround filters.

For the power-spectrum (complex-cell) responses, we computed the power spectrum of a large number of patches randomly sampled from natural textures. For each natural texture we found that the power at randomly sampled frequencies and orientations is, to close approximation, exponentially distributed and statistically independent. We used this approximation as the prior probability distribution for the power-spectrum responses.

#### Task-specific likelihoods

Once the input features are specified, and the priors measured, the task-specific features need to be specified. The task is to decide whether the two small patches are sampled from the same or different texture sheets, where the texture sheets are natural but arbitrary and unknown. For the task in Fig. 3A, there are two kinds of features available, the features created at the border between the two regions and the content features within two regions. For the task in Fig. 3B there are no useful border features, only content features.

We note up front that the proposed task-specific features are mathematically principled and were picked to extract as much information as possible from the small texture patches. They are not likely to be an accurate description of the task-specific features used by the visual system. However, evolution is likely to find features that capture similar information and we will see that human performance largely confirms this hypothesis.

Consider first the content features which are common to both tasks. Because the texture sheets are arbitrary and unknown, the Bayesian task-specific features are likelihood ratios (or equivalently log likelihood ratios) that the observed input feature responses from the two patches are from the same or from different probability distributions. The likelihood ratios are estimated by assuming that the feature responses are sampled from some family of probability distributions. Then, the parameters of the assumed family of distributions are estimated (by maximum likelihood) under the assumption that the parameters for the two patches are the same and that they are different. The likelihood ratio is the ratio of the two maximum likelihoods. We explored a number of possible families of probability distributions but discovered that many of the more sophisticated ones do not work well because of the limited data available in the 1×1 deg patches. In other words, the task requires one-shot learning of the probability distributions from the two small patches of texture. This strong limitation should also hold for local processing in visual cortex.

For the color and spatial-filter responses we found that good performance is obtained by assuming multinomial probability distributions for the single orthogonalized dimensions and then combining the log-likelihood ratios for the various dimensions. We use multinomial distributions because they can represent an arbitrary discrete probability distribution along any single axis. Estimating the parameters of the multinomial distributions requires specifying histogram bins for each of the orthogonalized dimensions. If the bins are either too coarse or too fine, the same-different discrimination performance declines. We find the optimal bins using a modified form of efficient coding which we call “adaptive histogram equalization” that uses the task-independent priors (more details in the Appendix). The log likelihood ratios (the task specific features) for each feature dimension are given by the following formula (derived in the Appendix):

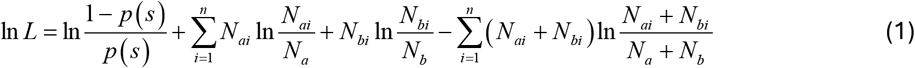

where *N*_*ai*_ is the number of response values in bin *i* for patch *a, N*_*bi*_ is the number of values for patch *b, N*_*a*_ and *N*_*b*_ the total number of values for patches *a* and *b, p* (*s*) is the prior probability the patches are from the same texture sheet, and *n* is the number of bins.

For the power-spectrum responses, the log likelihood ratio is given by the following formula (also derived in the Appendix):

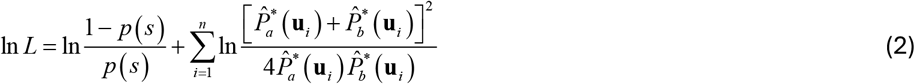

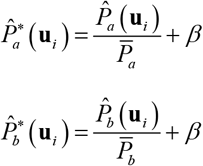

where 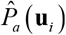 and 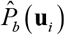 are the observed responses (power) in the two patches at the 2D frequency **u**_*i*_ = (*u*_*i*_, *v*_*i*_), 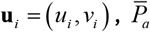 and 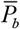 are the observed average responses in the two patches, and *β* is a fixed weak-power (noise) suppression constant picked to maximize performance. (Note that a 2D frequency is equivalent to a specific spatial frequency and orientation.)

To represent the effects of retinal eccentricity, we down-sample the input patches by the ratio of the midget-ganglion cell spacing at the given retinal eccentricity to that in the fovea, which is a down-sampling similar to that in primary visual cortex (see Appendix for details).

To train the model we first maximize performance for each task-specific feature (likelihood ratio). For the color and spatial-filter features this involves learning the bin bounds that maximize accuracy using adaptive histogram equalization. For the power-spectrum features this involves learning the value of *β* that maximizes accuracy. This learning specifies the calculation of all the specific log-likelihood ratios. We then learn the joint log-likelihood-ratio calculation for the content features using classification software we developed that assumes arbitrary multivariate normal distributions (Das & Geisler 2021).

Now consider the border features which are unique to the task in Fig. 3A. Here we consider just two features. One is the ratio of the total gradient energy perpendicular and parallel to the potential border:

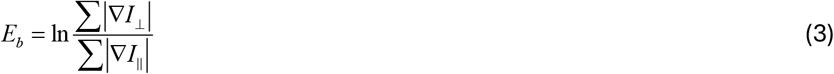

The other is the average of these same ratios along all lines away from the potential border but parallel to the potential border, 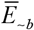. The gradient magnitudes were computed as above using the 1^st^ derivative steerable filter. The log likelihood ratio of the combined border features was again learned using our classification software (Das & Geisler 2021), as was the log likelihood ratio for the combined border and content log-likelihood ratios.

#### Predictions for same-different discrimination

We used two kinds of training stimuli (see thumbnails in the Appendix). One is a set of 60 Brodatz natural texture sheets. The second is a set of 60 Fabric texture sheets. We chose these manufactured Fabric textures, because they were generated in a very different way from the Brodatz textures and hence provide a useful comparison set for testing the generality of the decision variables learned during training.

Training was done three times: once for the Brodatz textures, once for the Fabric textures, and once for the combined textures. The training was done separately for four different retinal eccentricities. We found that the optimal decision bounds (and decision variables) in the feature space varied with eccentricity, but at each eccentricity they were very similar for Brodatz, Fabric, and combined textures, suggesting that fixed decision bounds at each eccentricity work well for a wide range of natural textures. Indeed, in ongoing work we are developing a self-supervised method for learning optimal decision variables and bounds from patches randomly sampled from arbitrary natural images. The decision variables learned this way are also quite similar to those learned for the Brodatz and Fabric textures. Here we show and use the decision variables and bounds learned for the combined Brodatz and Fabric textures.

The training procedure went as follows. First, we estimated the optimal decision variable (log-likelihood ratio) separately for each kind of content feature (power, histogram, and edge) in the task where the patches are separated (Fig. 3A). The learned decision variables were based on 36,000 randomly sampled patches from same and different textures (72,000 total patch pairs).

Second, we estimated the optimal decision variable in the three-dimensional content space where the axes are the same-different log-likelihood ratios for the three kinds of content features. The samples and decision bound (gray surface) that maximizes accuracy for foveal stimuli are shown in Fig. 6A. The accuracy with this decision variable and bound for Brodatz and Fabric patches is 93.6%. The axes represent the log-likelihood ratios for the three kinds of content features. The overall log-likelihood ratio that defines this decision bound is a principled measure of the content similarity between separated pairs of natural texture patches.

**Figure 6.**
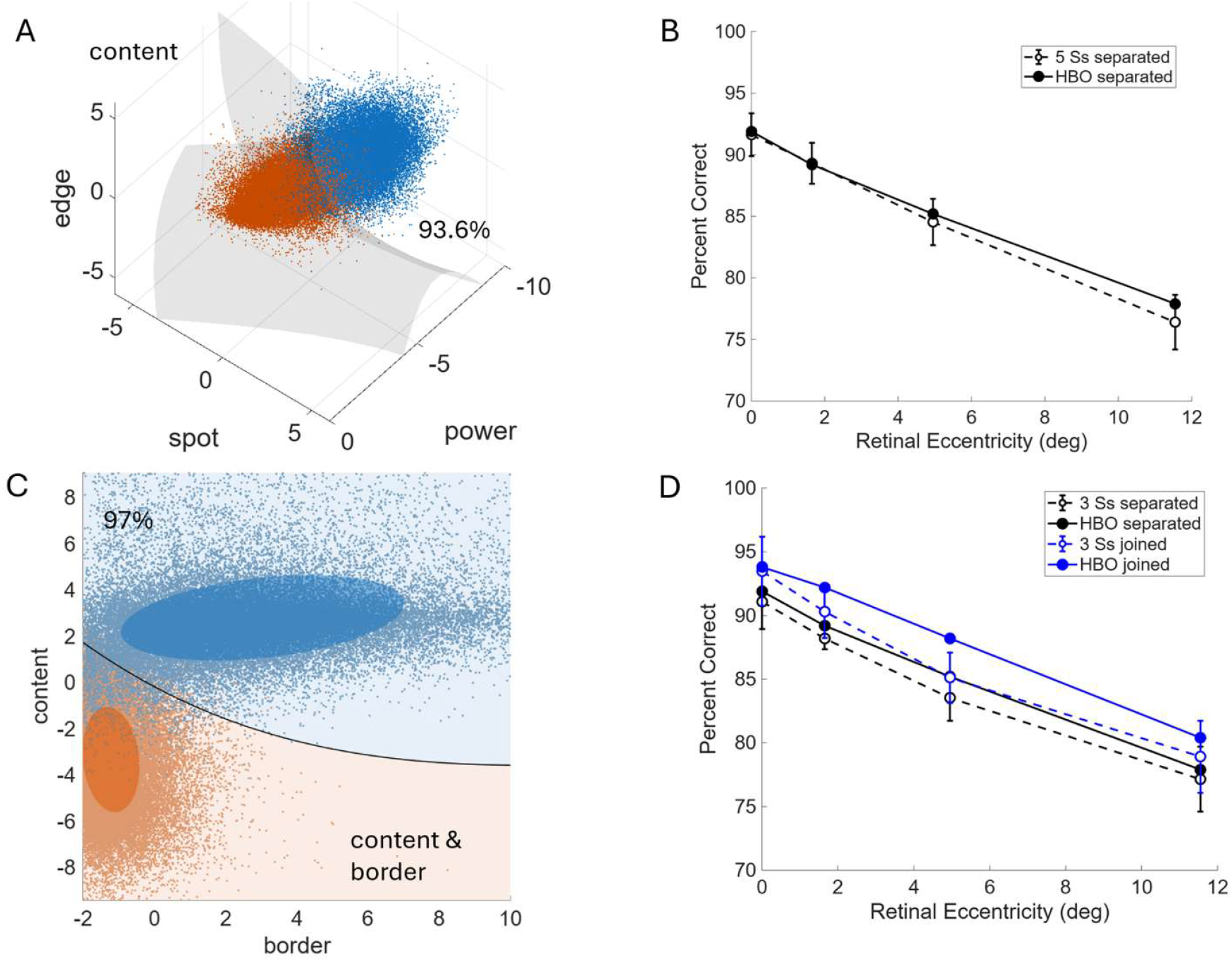
Discrimination results and predictions. **A**. Estimated optimal decision bound in the fovea, for same-different discrimination of texture patches in the separated condition, where two small texture patches are spatially separated and randomly sampled from the same or different Brodatz and Fabric texture sheets. **B**. The open symbols are the average same-different discrimination accuracy of five observers in the separated task as a function of retinal eccentricity. The patches were randomly sampled from the Brodatz texture sheets. (Error bars are standard deviations across observers.) The solid symbols are the predictions of the HBO discrimination model. **C**. The black curve is the optimal decision bound in the fovea for the same-different discrimination in the joined task for Brodatz and Fabric textures. Each blue dot is a pair of patches from a single texture, each red dot from different textures. The log-likelihood ratio associated with the decision bound is the measure of patch similarity. **D**. The open blue symbols are the average thresholds in the joined task and the open black symbols in the separated task for three of the five observers. The solid symbols are the predictions of the HBO discrimination model.

The solid black symbols in Fig. 6B show the accuracy achieved by the optimal decision bound for the four retinal eccentricities, when applied to only the Brodatz texture patches. The foveal accuracy is 92% correct. The estimated decision-bounds and accuracy levels are very stable. Freezing the bounds and testing with new randomly generated stimuli gave essentially identical results.

Third, we computed the content decision variable for the case where the texture patches are joined. Recall that in this case, when patches are “same”, they are really a single vertical rectangular patch of texture with a height that is twice the width; when “different”, they are random patches from different texture sheets. The accuracy for Brodatz and Fabric patches increased to 94.6% correct (93.3% for Brodatz alone) because joined patches that are “same” are more similar than random patches from the same texture sheet. Fourth, we learned the decision variable for the border cues. In this case, the accuracy was 87.1% for Brodatz and Fabric (83.9% for Brodatz alone). Finally, we learned the global decision variable for joined patches. The black curve in Fig. 6C shows the optimal decision bound which gives an accuracy of 96.8% correct (the axes are the values of the border and content log-likelihood ratios/decision variables). The solid blue symbols in Fig. 6D show the accuracy of the optimal decision bound for the four retinal eccentricities, when applied to only the Brodatz texture patches.

### Same-different Texture Discrimination Experiments

To test the HBO model of texture discrimination, human performance was measured for the two tasks shown in Fig. 3.

#### Methods

Same-different discrimination accuracy was measured as a function of retinal eccentricity. Stimuli were displayed on GDM-FW900 cathode-ray tube (CRT) monitor (Sony, Tokyo, Japan), with a size of 1920 × 1200 pixels, a refresh rate of 85 Hz, and a bit depth of 8. The stimuli were gamma-compressed prior to display on the screen. The stimuli were generated with MATLAB 2023a and the Psychophysics Toolbox (Brainard, 1997; Pelli, 1997).

On each trial, observers fixated a small gray square on a fixed luminance uniform background at 75 cd/m^2^. A pair of texture patches was then presented for 200 msec. Observers indicated with a button press whether the patches were sampled from the same or different texture sheets. The 64 × 64-pixel patches were randomly sampled from the 640 × 640-pixel Brodatz sheets (thumbnails shown in Appendix Fig. A4) and displayed at 60 pixels/deg. Each texture sheet was normalized to the mean luminance of 75 cd/m^2^ and an RMS contrast of 25%. The observer had unlimited time to respond, and the next trial began 1000 msec after the response. Auditory feedback was provided after each response.

Each session contained four blocks of 300 trials, one block for each of the four retinal eccentricities: 0°, 1.65°, 4.95°, and 11.55° (1200 trials per session). The pair of patches was always presented in the center of the screen. Observers used a chin rest and fixation was always straight ahead; specifically, the fixation point was moved on the display monitor, and the monitor was physically moved to keep fixation straight ahead. Each observer completed four sessions of the separated and/or joined conditions. Two observers did only the separated conditions. Three observers also did the joined conditions. When observers did both conditions, the two conditions were alternated across the eight sessions. The texture patches were randomly sampled on every trial and hence no two stimuli were presented twice to the same observer; however, all five observers saw the same stimuli.

#### Results

The open symbols in Fig. 6B show the average accuracy of the five observers in the separated task as a function of retinal eccentricity. As can be seen, the average accuracy of the observers is quite similar to that of the HBO model. This is a rather surprising result, because there was no fitting to the human-observer data. The features and the few parameters in the HBO discrimination model were adjusted only to maximize texture discrimination performance at each eccentricity and the predictions were not scaled overall to match human performance. The open symbols in Fig. 6D show the average accuracy of the three observers in the joined task. The average accuracy in the fovea of the HBO model is similar to average accuracy of the human observers. However, the HBO model performs better in the periphery, although the human observers do perform better in the joined task than the separated task. The higher accuracy of the HBO model suggests that the reduced sampling resolution of the periphery (in retina and V1) cannot fully explain the reduced performance in the joined condition. A potential explanation is the increase in intrinsic position uncertainty with retinal eccentricity (see Discussion).

Deeper insight into the texture discrimination data can be obtained by considering trial-by-trial correlations. An unbiased method for measuring trial-by-trial correlation is to measure decision-variable correlations (Sebastian & Geisler 2018; Qian et al. 2025). The advantage of decision-variable correlations (DVCs) over other methods is that they are based on signal detection theory and hence properly take into account the observer’s sensitivity (*d* ′) and decision criterion/bias. We computed DVCs to estimate the trial-by-trial correlation between the decisions of the different human observers, and between the decisions of the human observers and the model observer. The software returns the sensitivities, decision criteria, and decision-variable correlations. Fig. 7 plots the average DVCs across all pairings of the human observers, and the average DVCs between the human observers and the model observer.

**Figure 7.**
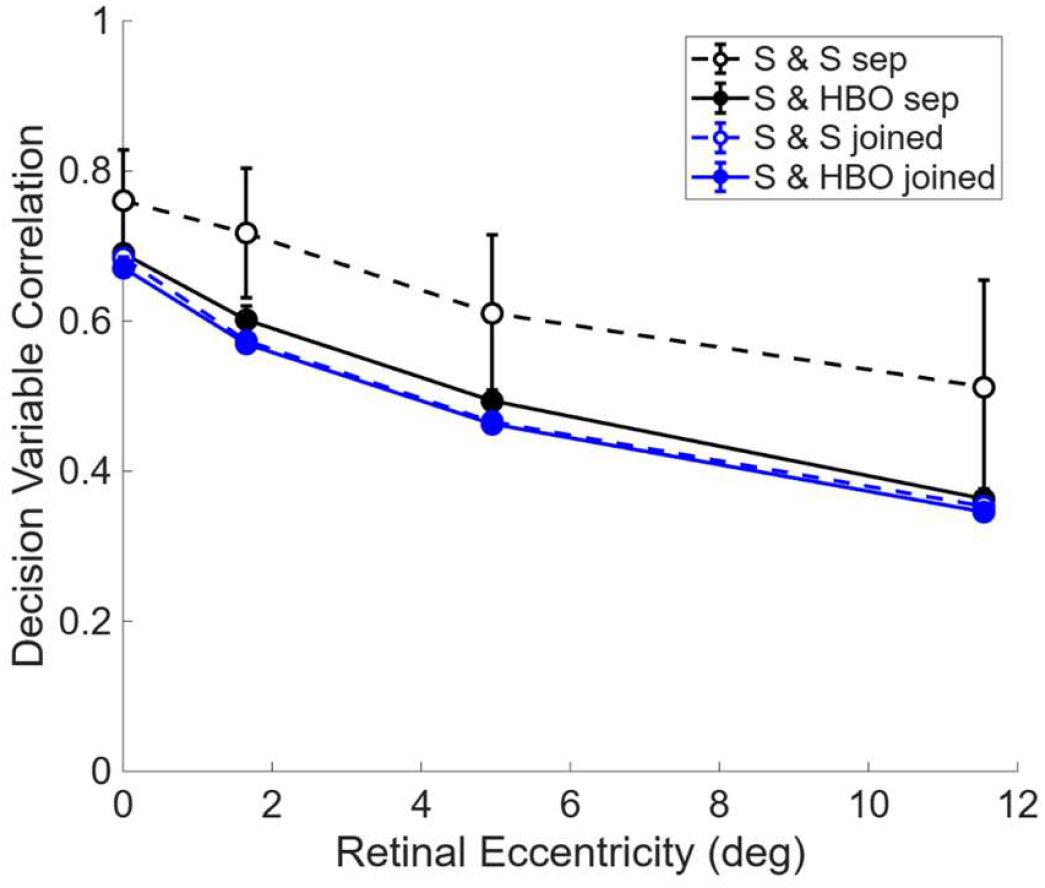
Average trial-by-trial decision variable correlations as function of retinal eccentricity. (Error bars are standard deviations across observers.)

Interestingly, the DVCs are quite high and decline modestly into the periphery. The high correlations between human observers suggest that much of the trial-by-trial variability is due to the variability of the stimuli, not due to neural variability, and that the different observers are using similar neural computations. The correlations are lower between the humans and model observer in the separated task, but still quite strong, especially in the fovea, suggesting that the humans and HBO model are using similar feature information. In the joined task, the DVCs between the human observers and between the human observers and HBO model are similar. Thus, the HBO model predicts trial-by-trial responses in the joined task as well as the responses of another human observer.

## Results: Texture Segmentation

The previous results section on texture discrimination focused on the two simple same-different discrimination tasks in Fig. 3. Here we step up to the bigger, more complex task of texture segmentation. We found, in the previous section, that the current HBO discrimination model approximately matches human performance for joined patches in the fovea and for separated patches in the fovea and periphery. Thus, the HBO model for local similarity grouping can serve as an appropriate input for evaluating the adequacy of the other grouping mechanisms of texture segmentation listed in Fig. 2, at least for the texture segmentation task where observers have time to fixate around the image. Thus, we now consider texture segmentation where foveal resolution is applied throughout the image.

### Grown Texture Region Images

As mentioned earlier, the goal is to understand the foundational texture segmentation mechanisms that do not depend on recognition of familiar objects or familiar scene context. To do this we designed a new kind of stimulus: grown texture region (GTR) images. These stimuli contain much of the complexity of natural images without including familiar objects or scene contexts. They allow controlled experiments and unlimited numbers of stimuli for training and testing models.

GTR images are generated using random Markov processes with the following steps: (1) select (possibly randomly) the number of texture regions, (2) randomly select seed locations for the texture regions to begin growing, (3) generate a random order of regions for the next growth pass, (4) for the current region, randomly select a region boundary location (boundary block), (5) for the current boundary location, randomly pick from the available directions for growth (left, right, up, down) and add a patch, (6) repeat from step 3 until all regions are processed, (7) repeat from step 2 until the space (background) is filled or until some fraction of the background is filled, (8) randomly assign a texture to each region and/or to the background.

A very wide range of texture region shapes are created by this procedure, and they have many of the properties of the texture regions created in natural scenes by growth, erosion, and occlusion. Many parameters of the procedure can be varied: the randomly selected textures, the number of texture regions (which may also be randomly varied), the fraction of the background texture region covered by the grown texture regions, the growth-block size, and growth bias parameters (e.g., a vertical growth probability bias). Fig. 8 illustrates a few possible kinds of grown texture regions. Fig. 8A was generated with the same parameters as those in Fig. 1. The parameters for Fig. 8B and 8C were the same except for the number of regions. The parameters for Fig. 8D were the same as in Fig. 8A except for the growth block size. The parameters for Fig. 8E were the same as in Fig. 8D except there was a bias toward central locations for growth seeds and growth was stopped once 50% of the background was covered. The parameters in Fig. 8F were the same as in Fig. 8E except there was a bias for growth in the vertical direction.

**Figure 8.**
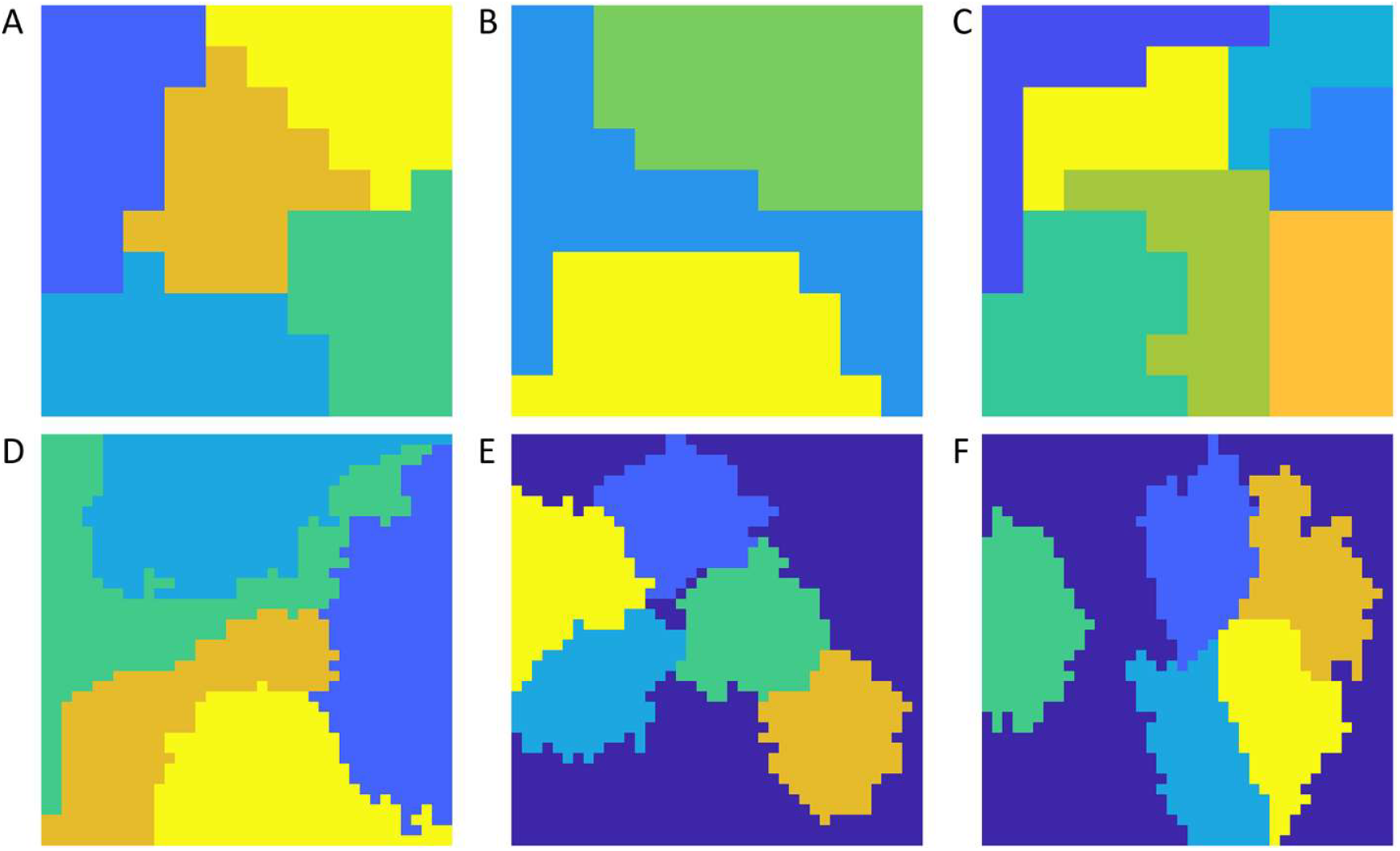
Examples of 640 × 640 grown texture regions. **A**. regions = 5, block size = 64, central bias = 0, coverage = 100%, orientation bias = 0 **B**. regions = 3, block size = 64, central bias = 0, coverage = 100%, orientation bias = 0. **C**. regions = 7, block size = 64, central bias = 0, coverage = 100%, orientation bias = 0. **D**. regions = 5, block size = 16, central bias = 0, coverage = 100%, orientation bias = 0. **E**. regions = 5, block size = 16, central bias = 80%, coverage = 50%, orientation bias = 0. **F**. regions = 5, block size = 16, central bias = 80%, coverage = 50%, orientation bias = vertical.

### Hierarchical Bayesian Observer Model for Texture Segmentation

#### Local similarity grouping

The first step of the HBO model of texture segmentation is to apply the above local similarity grouping model for the task in Fig. 3A to every pair of neighboring texture patches. Here we use only the HBO model for the fovea and thus we are assuming that the segmentation is obtained by fixating around the image. For most patches there are four log likelihood ratios (based on content and border cues) from the patches that surround it. The log likelihood ratios can be regarded as similarity measures, the larger the log likelihood ratio the greater the pairwise similarity. Let *φ*_*ij*_ be the log likelihood ratio (similarity) between two patches. When the patches are neighboring, the similarity is based on content and border cues (combined content and border similarity).

#### Mutual similarity grouping

In the next step, the content similarities are computed between all pairs of patches at all distances. The mutual similarity *µ*_*ij*_ between a neighboring pair of patches is defined to be the cosine similarity between the vector of all content similarities to patch *i* and the vector of all content similarities to patch *j* (the dot product divided by the product of the vector lengths):

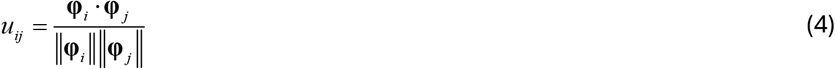

where the bold symbols represent the similarity vectors. Intuitively, if a neighboring pair has the same pattern of similarities to other patches, then they are more likely to be from the same texture region.

The combined similarity for each neighbor pair is given by

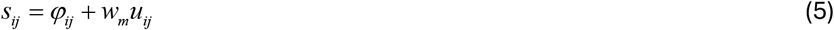

where *w*_*m*_ is a constant that is adjusted to maximize overall performance. Mutual similarity can be thought of as a form of recurrent global feedback to the local similarity between neighboring pairs of patches. The similarity of each neighboring pair *s*_*ij*_ is compared to a grouping criterion *γ* _*l*_. If the similarity exceeds the criterion, the pair is nominally labelled as same, and below the criterion as different. However, similarities that are too close to the criterion are regarded as uncertain. Specifically, if the distance from the criterion is less than some criterion distance, |*s*_*ij*_ – *γ* _*l*_ |< *γ* _*c*_, then the link is labelled as uncertain, and not used in the transitive grouping step. This suppression of weak links is a component of confidence grouping. The constants *γ* _*l*_ and *γ* _*c*_ are also adjusted to maximize performance.

#### Transitive grouping

The next step applies transitive grouping (a global grouping process) to all pairs labelled “same.” As mentioned earlier, if patch ***a*** groups with patch ***b***, and patch ***b*** groups with patch ***c***, then patches ***a*** and ***c*** are grouped. Previous applications of transitive grouping have been to contour elements (e.g., Geisler et al. 2001); however, the principle applies equally well to texture content. Transitive grouping creates an initial segmentation of the image into regions, although there are sometimes a few isolated regions that consist of only one patch. Transitive grouping is a crucial process that allows grouping of textures that are stationary as well as textures that are nonstationary but smoothly varying. In images of the natural environment, smoothly changing textures are common because of perspective geometry and physical processes that grow/create surfaces. Transitive grouping makes segmenting these textures possible. On the other hand, because it causes grouping to propagate it requires some control. Specifically, an accidentally strong similarity between a pair of patches in two large texture regions can cause the regions to merge into one region (see below).

#### Confidence grouping

Confidence grouping helps to control transitive grouping. Low confidence similarities (similarities close to the same-different decision bound) are places where transitive grouping could inappropriately merge regions together, thus they are ignored in the transitive grouping step. For many low-confidence links, the two patches still end up in regions following transitive grouping, because there are generally four similarities computed for each patch (left, right, up, and down) and any one of those could pull the patch into a region. For any patch that is not in any region (a rare event), its content similarity is computed for all patches in the segmented regions that they touch. They are then assigned to the region with the highest average similarity.

#### Region similarity grouping

This compares average content similarity of the regions obtained with the first four steps. If the average content similarity of two regions exceeds a criterion *γ*_*r*_, then the regions are merged. Occasionally, a region is isolated in that it consists of just one patch. That patch is merged with whatever neighboring region best matches it.

#### Segmentation Simulations

The earlier section on texture discrimination described our efforts to develop an HBO model of same-different discrimination for 1° x 1° texture patches randomly selected from natural texture sheets. The model combined measurements of natural image statistics and biologically plausible content and border features, with efficient coding and Bayesian decision-making. The few parameters in that model were adjusted to maximize discrimination performance for a wide range of textures. Interestingly, the model does a reasonable job of predicting human discrimination performance on Brodatz textures, even though it was not trained to predict human performance. Thus, the local grouping component of our HBO model for texture segmentation is consistent with human local texture discrimination performance. We do not yet have measurements of human segmentation performance for GTR images, but we can ask how well the proposed global grouping mechanisms work given local grouping mechanisms that are consistent with human performance. To do this we simulated texture segmentation of GTR images containing Brodatz and Fabric texture (thumbnails of the texture sheets are in the Appendix).

There are four parameters in the texture segmentation model: the scalar, *w*_*m*_, controlling the contribution of mutual similarity to the local similarity, the criteria, *γ* _*l*_ and *γ*_*c*_, controlling the grouping of neighboring patches, and the criterion, *γ* _*r*_, controlling the region grouping. We systematically, but coarsely, varied the four parameters to maximize segmentation accuracy. There are multiple combinations of the four parameters that achieve similar accuracy. The parameter values were fixed at one of these combinations for all the simulation results shown here (see caption of Fig. 9).

**Figure 9.**
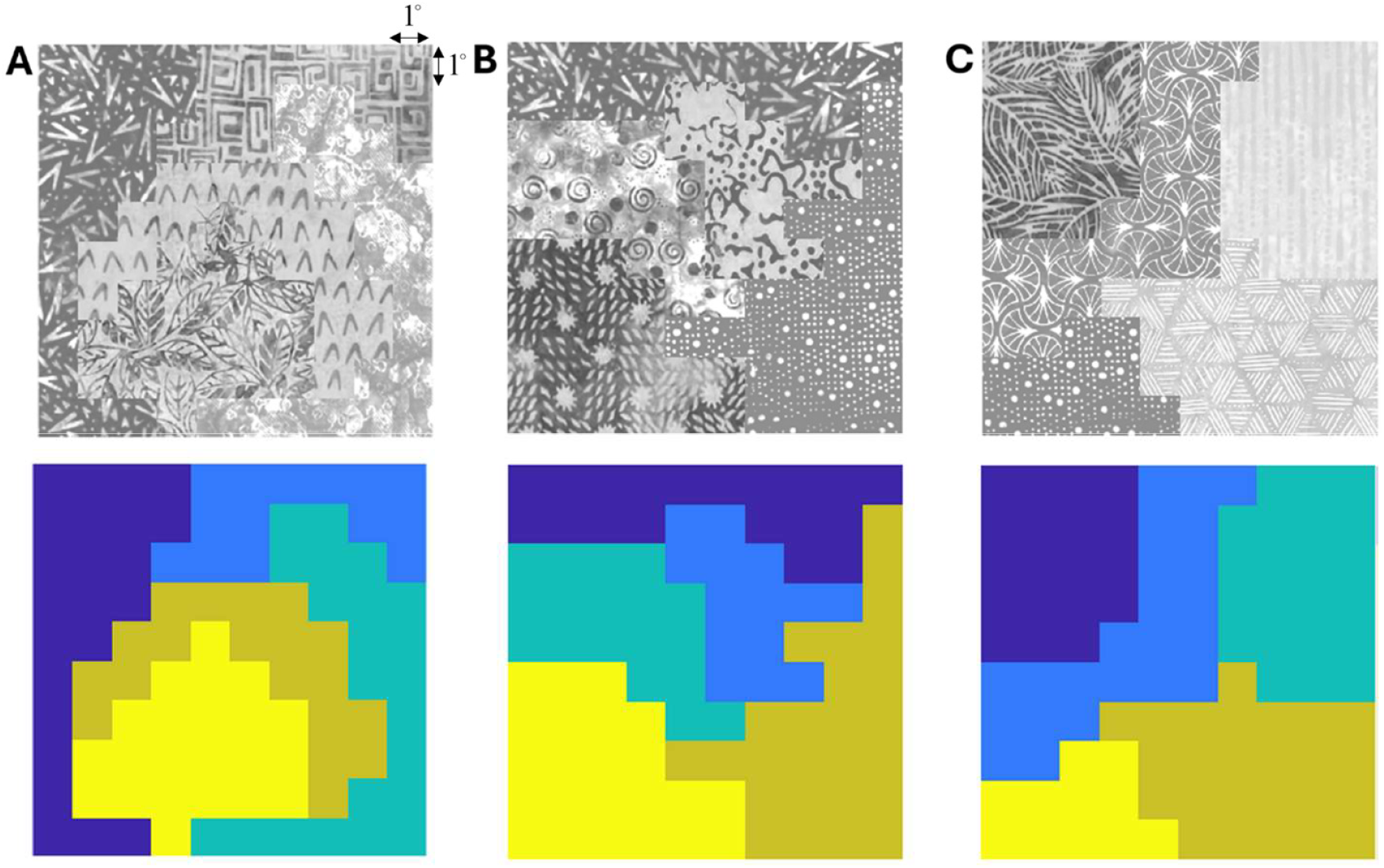
Examples of perfect segmentation of grown texture region images for Fabric textures. See Fig. 1 for examples for Brodatz textures. HBO segmentation model parameter values: *w*_*m*_ = 7, *γ* _*l*_ = 6, *γ* _*c*_ = 1.0, *γ* _*r*_ = 1.5

The HBO model worked surprisingly well, given the heterogeneity and non-stationarity of many of the textures. There are many instances where the segmentation of the entire image was perfect. Three examples for the Brodatz textures are shown in Fig. 1 and three examples for the Fabric textures are shown in Fig. 9.

Approximately 54% of all Brodatz GTR images and 46% of all Fabric GTR images were segmented perfectly, and 76.5% of Brodatz GTR regions and 72% of Fabric GRT regions were segmented perfectly (see Fig. 10). This level of success is largely (but not solely) due to the local similarity and transitive grouping.

**Figure 10.**
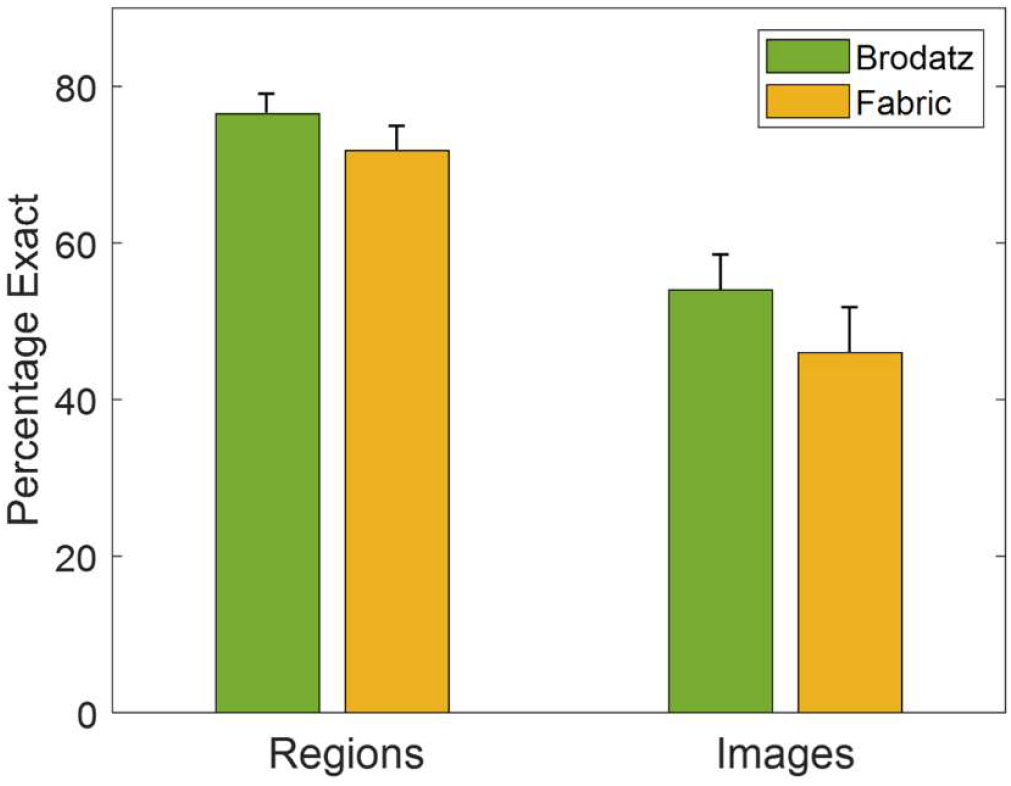
Left: percentage of texture regions segmented perfectly. Right: percentage of GTR images segmented perfectly. (Standard errors are based on 600 regions and 120 images.)

The failures of segmentation tend to be of three different types (Fig. 11). One kind of failure is when the textures are so similar that border and content cues in the two regions are relatively weak (Fig. 11A). This is a straightforward failure that is likely to be consistent with failures of the human visual system.

**Figure 11.**
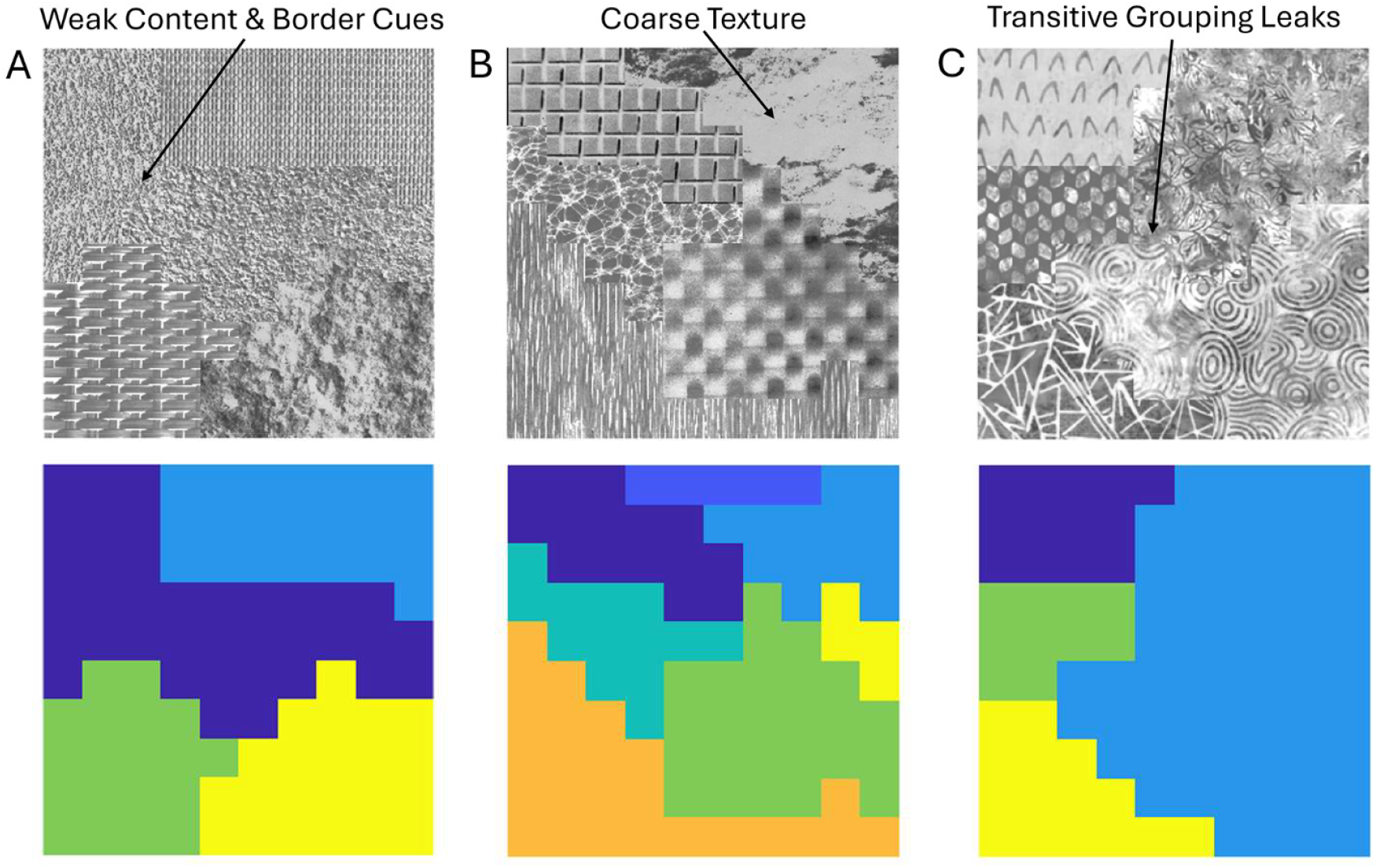
Typical reasons for failure of perfect segmentation. **A**. Brodatz GTR image with weak content and border cues between two ground-truth regions. **B**. Brodatz GTR image with very coarse texture in one region. **C**. Fabric GTR image with accidently weak content and border cues at one location along border between two ground truth regions. Number of perfectly segmented texture regions for images in A, B and C are 3, 4 and 3, respectively.

A second kind of failure seems to be due to the coarseness of the texture in the region. Some of the textures are so coarse that they get divided into multiple regions. For example, the upper right region in Fig. 11B gets partitioned into three regions. The three regions are not unreasonable at a small scale, but most human visual systems probably see the region as filled with a coarse material of some sort. This is the kind of region that might be more accurately segmented with HBO models that consider a wider range of scales (e.g., spatial frequencies less than 1 c/deg).

The third kind of failure appears to be due to leaks in the transitive grouping process (Fig. 11C). These occur when there is an accidentally high similarity between neighboring patches from two different kinds of texture, even though the texture patches in the two regions are generally quite different. Because of transitive grouping the two regions get bound together. Confidence grouping helps reduce this kind of failure but does not always work.

In sum, these simulations of the HBO model of segmentation suggest that the relatively simple, but principled, grouping mechanisms described here may be sufficient to account for much of human ability to segment scenes that do not contain recognizable objects or context.

## Discussion

The aim of this study was to gain a better understanding of texture segmentation under naturalistic conditions, for the case where the images do not contain recognizable objects, materials, or scene context. We argue that the mechanisms supporting segmentation under these conditions are the backbone for the more typical case where images do contain recognizable objects, materials, and context. Specifically, recognition-free mechanisms are required for recognition learning, and after the learning they provide the initial segmentations that help drive the recognition mechanisms.

We described a relatively simple hierarchical Bayesian observer (HBO) model of texture segmentation with five grouping mechanisms: local-similarity grouping, mutual-similarity grouping, transitive grouping, confidence grouping, and region-similarity grouping. We first found that a principled HBO model of texture discrimination predicted human same-different discrimination accuracy for small, separated texture patches as a function of retinal eccentricity and for joined texture patches in the fovea. However, humans did not quite reach model performance for joined patches in the periphery. The HBO model’s predictions were essentially parameter free, because the features and the few parameters were adjusted to maximize the model’s performance and not to fit the human performance. The combined border and content decision variable is the measure of local similarity. We then asked whether this local similarity grouping mechanism, which is consistent with human discrimination performance in the fovea, would be sufficient to support segmentation of images having random shaped regions filled with random textures (GTR images). We found that the HBO model, with its four additional grouping mechanisms, did a very reasonable job of segmenting both Brodatz and Fabric GTR images.

Many of the Brodatz and Fabric textures are non-stationary, with their statistical properties varying over space. The success with these non-stationary textures is largely due to transitive grouping. In the same way that transitive grouping can link together smoothly changing orientations of contour elements (e.g., Geisler et al. 2001), it can link together smoothly changing texture properties. Many of the Brodatz and Fabric texture regions that were accurately segmented contain textures with properties that vary smoothly over space.

### Transitive grouping and 3D surface geometry

Even statistically stationary 3D surface textures often give rise to nonstationary image textures because of perspective projection. Thus, because of transitive grouping, it is possible that the images of 3D surfaces can be correctly segmented even by HBO models trained only on fronto-parallel (2D) textures. As a quick test, we created GTR images from natural images of slanted planar surfaces. Two examples of these are shown at the top of Fig. 12. Even though the scale and aspect ratio of features within the texture regions changes, the HBO model segments them reasonably well. The only error here is in the upper right corner of the image in Fig. 12B.

**Figure 12.**
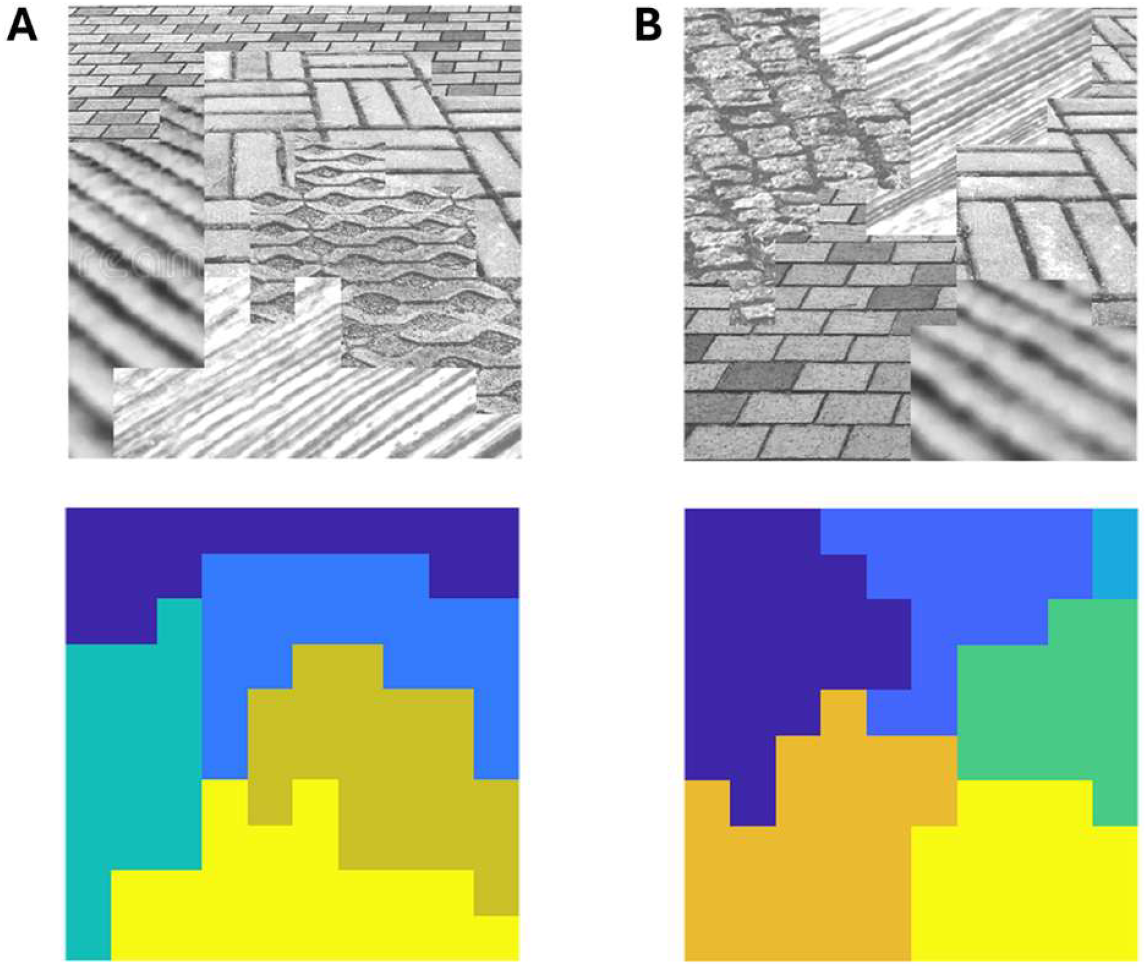
Segmentation of texture regions that are images of surfaces slanted in depth. This demonstrates the effectiveness and importance of transitive grouping. Note the segmentation error in the upper right corner of example B. This failure occurred because region grouping failed (the patches in the small region are not very similar to any of those in the large region). If the texture regions are correctly segmented, then the systematic changes in the texture can be used to estimate 3D geometry.

Accurate segmentation of texture regions created by perspective projection could play an important role in recovering 3D geometry. Specifically, once a region has been accurately segmented, the variation of feature scale and aspect ratio within the region can be used to help infer local surface slant and tilt within the region. One consequence of this inference is the creation of shape illusions. For many regions in Fig. 12 the shape of the region looks quite different in the upper and lower images. For example, compare the apparent shape of the gold region in the bottom image of Fig. 12A with the corresponding region in the upper image.

These preliminary results suggest that investigation of HBO models of segmentation for slanted and curved surfaces may be a fruitful direction for further study.

### Normalization: texture segmentation and illumination estimation

Luminance and contrast normalization probably play an important role in both texture segmentation and in the estimation of scene illumination. The HBO model performs local luminance and contrast normalization before computing local similarity. This step can allow accurate segmentation of texture regions even if the illumination changes over the scene. This is demonstrated in Fig. 13 A and B. Specifically, the GTR image in Fig. 1A was regarded as a reflectance image and the image in Fig. 13A shows a simulation of the effect of modulating the illumination across the upper and lower halves of the image. Fig. 13B shows that the segmentation of the texture regions remains accurate. Fig. 13C shows the map of all the luminance normalization scalars. If these signals are also properly processed, they could provide an estimate of the illumination. In other words, by using both the locally normalized texture images and normalization signals it is possible to simultaneously segment the texture regions and estimate the approximate pattern of illumination. Contrast normalization probably also plays an important role in allowing segmentation when the direction of lighting changes, because lighting direction modulates the contrast of non-smooth surfaces. This is yet another important benefit of luminance and contrast normalization mechanisms, which are known to have many other benefits (Albrecht & Geisler 1991; Heeger 1991; Geisler & Albrecht 1997; Carandini & Heeger 2012; Zhang & Geisler 2025). These preliminary results suggest that HBO texture segmentation models could serve as the foundation for principled models of illumination estimation in natural scenes.

**Figure 13.**
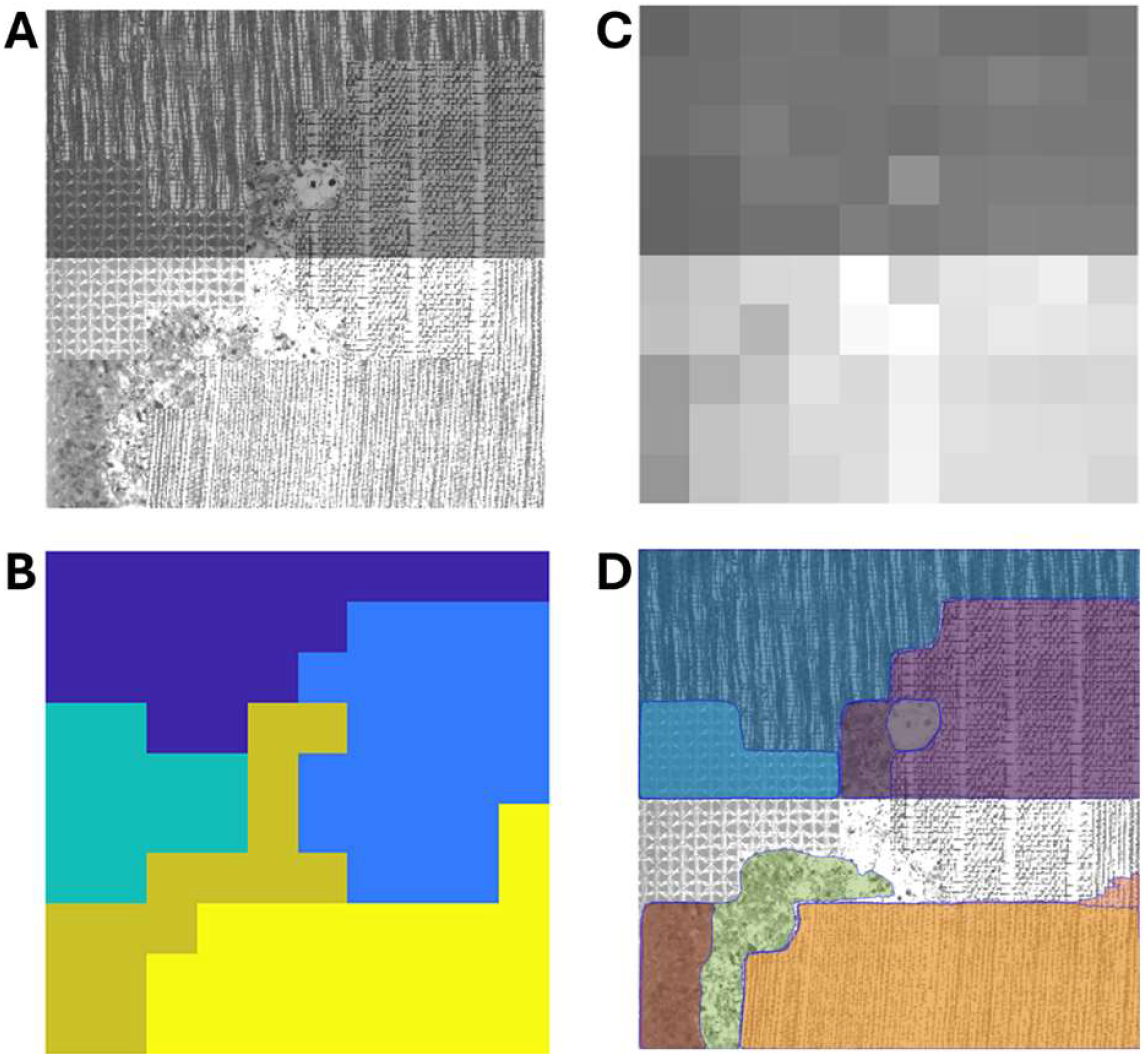
Estimation of illumination from local luminance normalization signals. **A**. GTR image from Fig. 1A but with a simulated step change in illumination. **B**. Normalization allows accurate segmentation of texture regions even when illumination is nonuniform. **C**. The normalization signals shown here can then be used to estimate the illumination. **D**. Meta’s Segment Anything Model (SAM) does not have something equivalent to local normalization and hence cannot take illumination into account, leading to relatively poor texture segmentation.

### Current state-of-the-art image segmentation

At the time of writing this report, the state of the art in image segmentation is very large-scale machine learning algorithms that combine large-language models and computer vision. An excellent example is Meta’s impressive “segment anything model (SAM)”, which was trained on a billion image-region masks using a mixture of unsupervised and supervised training (Kirillov et al. 2023). SAM is designed as a foundation model that can adapt to a wide range of tasks and that makes use of all kinds of useful low-level and high-level information for segmenting natural images. It was not trained on GTR images which contain little or no high-level information. Nonetheless, it is informative to test whether it can segment GTR images in a fashion consistent with humans and with our HBO models. Fig. 14 shows SAM’s segmentations of the GTR images in Figs. 1 and 8. It finds some sensible sections of region boundary and segments some of the regions correctly; for example, the upper gray, gold, and purple regions within the three images in Fig. 14A. However, it tends to split up the ground-truth regions into many subregions. SAM does hierarchical segmentation, so in some cases this is not inappropriate, in that humans might agree with the hierarchy. But in many cases, humans would never split the region the way SAM does. For example, humans would never segment the upside-down V texture region of the first image in Fig. 14B into different regions, or the vertical contour texture starting in the lower left of the middle image in Fig. 14A.

**Figure 14.**
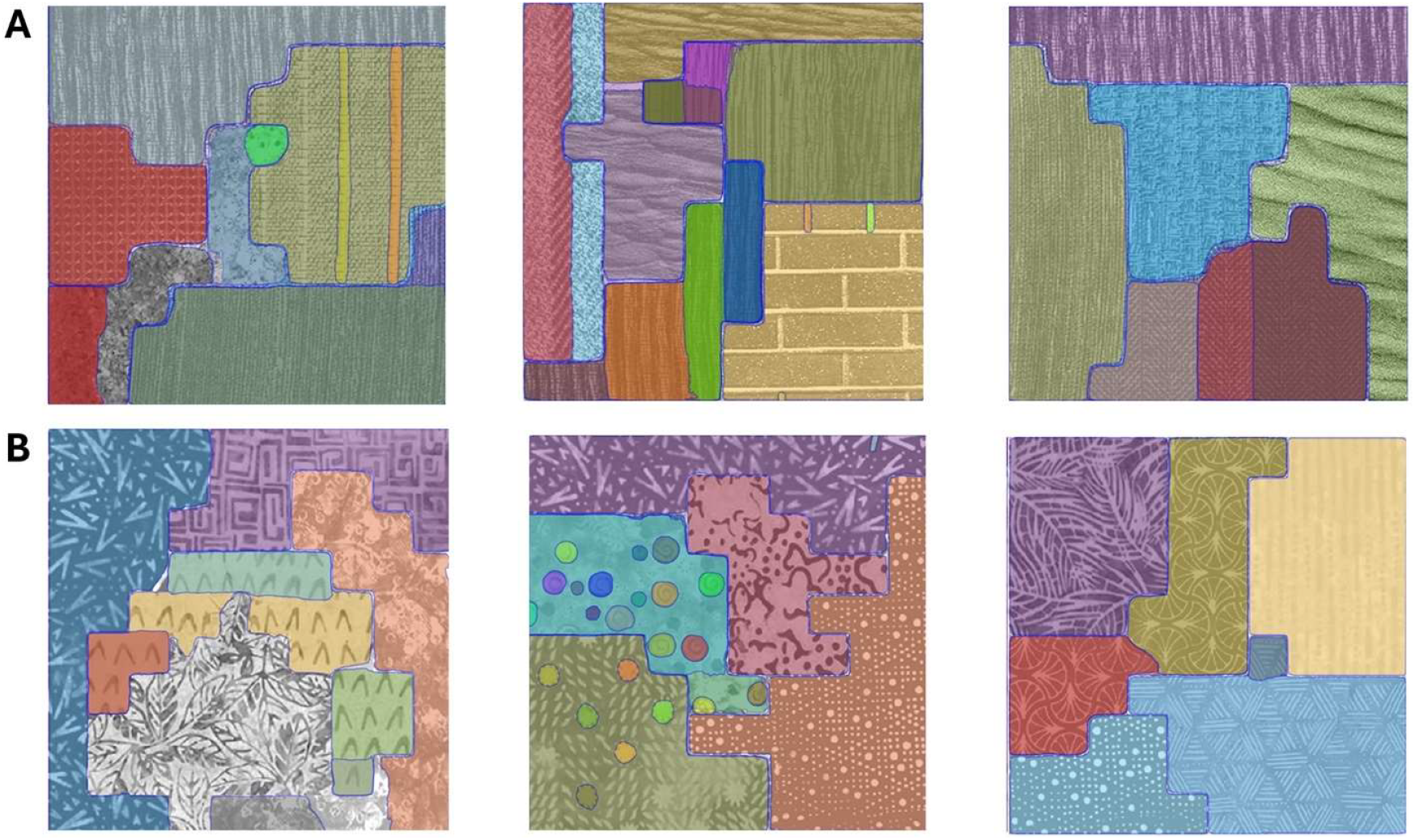
Segmentations of GTR images by Meta’s Segmentation Anything Model (SAM). **A**. Brodatz GTR images in Fig.1. **B**. Fabric GTR images in Fig. 8. SAM often splits regions in ways that the human visual system does not. See also Fig. 13D.

Unlike the human visual system, SAM has no explicit luminance and contrast normalization stage and hence has even more trouble when there is variable illumination. As can be seen in Fig. 13D, SAM now makes more severe segmentation errors. For example, it groups four different kinds of texture into a single region (gray region in the middle of the GTR image). It is possible that explicit incorporation of the low-level Gestalt principle of continuity (transitive grouping), as well as local luminance and contrast normalization, could improve the robustness of machine learning algorithms and allow them to perform better when there are fewer recognizable objects and materials, and/or less context.

### Task-independent priors

A novel feature of the HBO models is the measurement and use of task-independent priors to constrain and train the task-dependent features. The task-independent priors are not guaranteed to improve performance from the task-dependent features, but they do not hurt performance, and we found that they did improve performance for most of the task-dependent features we considered. Our view is that measuring the priors is a generally sensible thing to do because it is easy, can improve the task-dependent features, and can provide additional insight into the structure of natural scenes. Determining the task-independent priors of well-characterized neurons in the early sensory systems of a species could be useful for generating hypotheses for subsequent circuits.

### Relative contribution of content and border features

What is the relative contribution of content and border information to discrimination of joined (neighboring) natural-texture-patch discrimination? To address this question, we compared performance of the two classes of discrimination cues as function of retinal eccentricity separately for Brodatz and Fabric textures. Recall that optimal decision bounds for border, content, and joint cues were trained on samples from all Brodatz and Fabric textures. We froze those decision bounds and then applied them separately to joined-task samples from Brodatz and Fabric textures.

The results are shown in Fig. 15. The solid symbols are results for the Brodatz textures and open symbols for Fabric textures. The black symbols show the performance for all cues. The solid black curve is the same as the solid blue curve in Fig. 6D. The overall performance for Brodatz and Fabric textures is similar with a slightly faster falloff with eccentricity for the Fabric textures. Accuracy for the content cues alone is given by the blue symbols and for the border cues alone by the red symbols. Overall, the border cues are less useful than the content cues and their usefulness decline faster with eccentricity. The border cues are more useful for the Fabric than Brodatz textures, especially in the fovea. The steeper falloff for border cues is perhaps not surprising because the useful border information is most likely contained in higher spatial frequencies. Comparison of the solid blue curve here with the solid black curve in Fig. 6B shows that the content cues in the joined condition gives better performance than the content cues in the separated condition. The fact that content cues work better for neighboring patches increases the usefulness of transitive grouping in the texture segmentation task.

**Figure 15.**
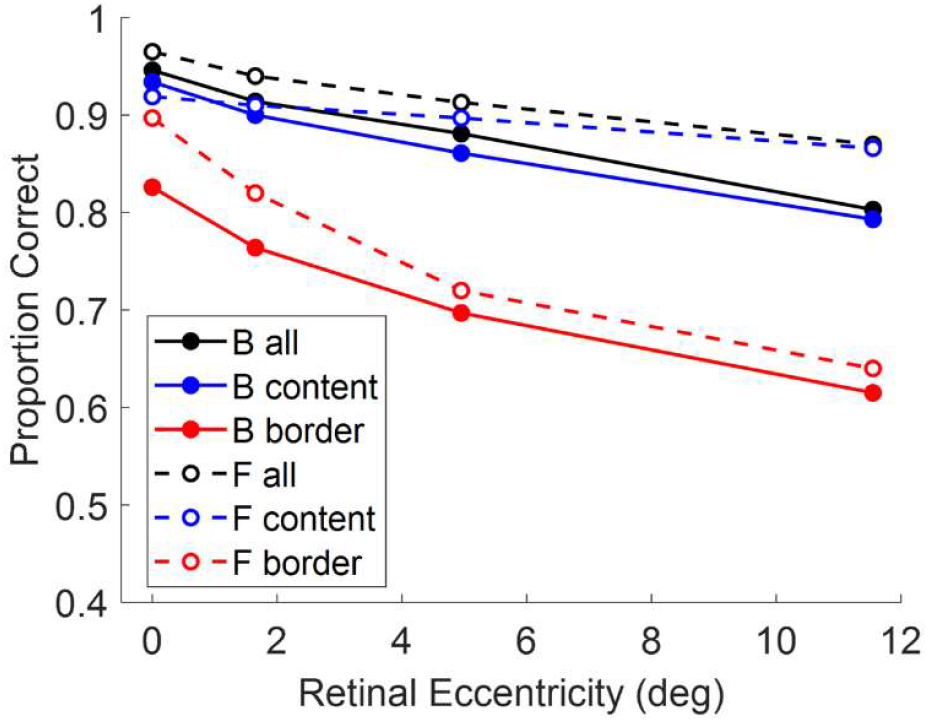
HBO model accuracy as a function of retinal eccentricity for all cues, content cues, and border cues, for Brodatz (B) and Fabric (F) texture patches in the joined task.

### Universal decision bound for local-similarity grouping

The black curve in Fig. 6C shows the optimal decision bound for foveal same-different discrimination in the joined task when trained with random patches of Brodatz and Fabric textures. The log-likelihood ratio (decision variable) associated with this bound is the measure of local similarity grouping strength used in the HBO model of texture segmentation. The gray surface in Fig. 6A shows the optimal decision bound for foveal same-different discrimination in the separated task when trained with random patches from Brodatz and Fabric textures. The log likelihood ratio (decision variable) associated with this bound is the measure of similarity grouping strength used for non-neighboring patches in the HBO model of texture segmentation.

We have also trained these decision variables and bounds with Brodatz textures alone and with Fabric textures alone. The bounds are very similar to the bounds in Fig. 6, and the model performance is very similar. For example, when trained on Brodatz and Fabric textures in the joined task, the foveal accuracy when tested with Brodatz textures is 93.8% correct and when tested with Fabric textures is 96.9%. When trained and tested on Brodatz textures the accuracy is 95.6%, and when trained and tested on Fabric textures the accuracy is 96.9% correct. Thus, the general bound and decision variable gives accuracies that are within a couple percent of those obtained with training on the same kind of texture. These results suggest that there may be a universal decision-bound that works well for a very wide range of natural textures. In preliminary work, we are using a self-supervised method to learn optimal decision variables and bounds for patches randomly sampled from natural images. Initial results show that these decision variables and bounds are similar to and work as well as those trained on Brodatz and Fabric textures. The existence of an approximately universal decision bound suggests that a relatively small, fixed set of low-level mechanisms could evolve to support efficient local-similarity grouping, especially in species with a shorter lifespan for learning and with fewer neural resources.

### Intrinsic variability

One puzzling result from the same-different texture discrimination experiments is that human performance fell below that of the HBO model in the periphery for the joined condition, but not for the separated condition and not for the joined condition in the fovea (see Fig. 6D). One possibility is that intrinsic position uncertainty and potentially other sources of internal variability increase with retinal eccentricity (Michel & Geisler 2011). This internal variability might be more damaging for the border cues which have a precise location. Also, content tends to change less for joined patches that are from the same texture than separated patches from the same texture. Thus, the log-likelihood ratio tends to be further from the decision boundary in the joined case. Intrinsic variability randomly moves some patches closer to the decision boundary, reducing performance. When the patches are separated, the effect of intrinsic variability is swamped by the variability of the of random patches from the same texture, and hence performance is not reduced. The fact that the decision-variable correlation between subjects and between the HBO model and the subjects is lower in the periphery (Fig. 7) is also consistent with an increase in intrinsic position uncertainty.

### Limitations and future directions

The emphasis here is on achromatic spatial pattern information. Chromatic information is also useful for texture segmentation under natural conditions. For example, including the chromatic information in the Fabric textures substantially improves performance because of the wide variety of color in the Fabric dataset. In most natural scenes chromatic information is on average weaker. An important future direction is to examine the role of chromatic information.

It is possible that better models of local similarity grouping could support better texture segmentation performance without changing or adding to the last four steps. The HBO discrimination model already reaches or exceeds human texture discrimination performance (Fig. 6) so it might be argued that better features for similarity grouping would not be plausible (although different but not better features would be plausible). However, we found that the between subject DVCs, while substantial, were still less than 1.0 (an average of 0.7 to 0.75 in the fovea). The amount of intrinsic variability needed to produce this trial-by-trial correlation (assuming the observers are using the same features) should allow some room for additional or better features in the human visual system. As mentioned earlier we tried a number of other features, but the more complex features failed to help because of the small one-degree patch size. Nonetheless, it is possible that machine learning tools such as variational autoencoders may be able to find features that improve local-similarity grouping.

Finally, another important next step will be devising texture segmentation tasks with GTR images that allow testing hypotheses about the steps subsequent to local-similarity grouping.

## Acknowledgements

Supported in part by NIH grants EY11747 and EY024622.

## Appendix

### Multinomial log likelihood ratio

Let **N**_*a*_ = (*N*_*a*1_, …, *N*_*an*_) and **N**_*b*_ = (*N*_*b*1_, …, *N*_*bn*_) be the vectors of observed counts in the *n* histogram bins for patches *a* and *b*. According to Bayes’ rule the posterior probability that the two patches are random samples from the same distribution is given by

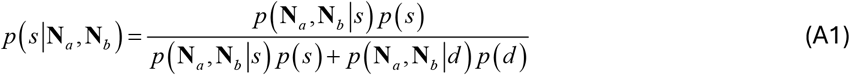

therefore, the log likelihood ratio is given by

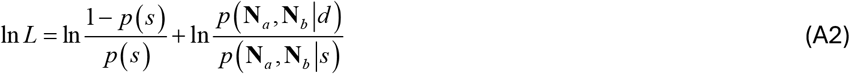

where *p* (*s*) is the prior probability that the patches are “same.” Assuming multinomial probability distributions the log likelihood ratio is given by

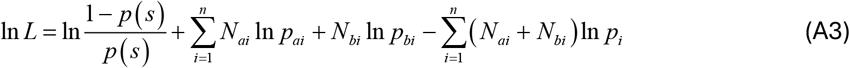

where *p*_*ai*_ and *p*_*bi*_ are the probabilities of observing a value in bin *i* if the distributions are different and *p*_*i*_ is the common probability if the distributions are the same. The maximum likelihood estimate of the three probabilities given the observed histograms are 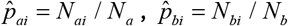 and 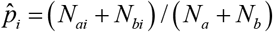, thus the empirical log likelihood ratio is

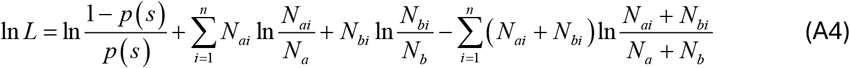

We assume that *p*_*ai*_ > 0, *p*_*bi*_ > 0, *p*_*i*_ > 0, thus (for example) if *N*_*ai*_ = 0 then *N*_*ai*_ ln 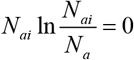.

### Adaptive histogram equalization (AHE)

Histogram equalization is a classic form of efficient coding that encodes random variables more finely in the regions where the random variable has higher probability (i.e., assigning more bits to ranges of values that occur more frequently). This is a principled strategy that maximizes the entropy of the encoding for a given number of bins (symbols). Although maximum entropy encoding of the prior feature distributions is not guaranteed to give optimal bins, it is a good place to start. We tried simple histogram equalization (HE) of the 9 CDFs in Figs. 5C-D, for different numbers of bins. For each number of bins, we measured accuracy for the same-different task in Fig. 3B for a large number of trials with Brodatz textures and Fabric textures. We found that accuracy was optimal for some intermediate number of bins and that the optimal number of bins varied with the specific feature. We used the performance for these bins as a benchmark for comparison with the more flexible AHE method described by the following steps.

Step 1: Find the bin bound that splits the probability distribution into two equal areas (e.g., the bin bound corresponding to the CDF probability of 0.5) and measure accuracy in the same-different task in Fig. 3B, for a large number of trials with Brodatz and Fabric textures.

Step 2: Find the bin bound that splits the first bin into equal areas (e.g., the bin bound at 0.25 on the CDF). Measure performance again on the same stimuli. If performance improves more than some small criterion fraction of the current error rate (0.4%) then keep the bin; otherwise freeze the bin.

Step 3: Find the bin bound that splits the next unfrozen bin into equal areas (e.g., the bin bound at 0.75 on the CDF). Measure performance again on the same stimuli and keep the split if performance improves more than the criterion; otherwise freeze the bin.

Step 4: Repeat until all unfrozen bins are checked in the current pass. If all bins are frozen stop; otherwise, go back to Step 2 and split the remaining unfrozen bins one at a time.

Fig. A1 shows the bins found in the fovea for the 9 feature dimensions. The optimal number of bins is indicted near the top of each plot. We obtained optimal bins for the fovea and for the three other retinal eccentricities in Fig. 6. Overall, the AHE bins gave better performance than the HE bins, but the improvement was modest, ranging from an increase of 0% to 4% correct, with a mean increase across all dimensions of ~1.3% correct.

Interestingly, almost all the optimal bins for the center-surround filters are in the negative range, suggesting that most of the useful texture information for these filters are in small dark spots. This seems consistent with the fact that there are more off-center than on-center neurons in visual cortex (Mazade et al. 2019) and with evidence for texture discrimination mechanisms tuned to black spots (Chubb et al. 2004).

**Figure A1.**
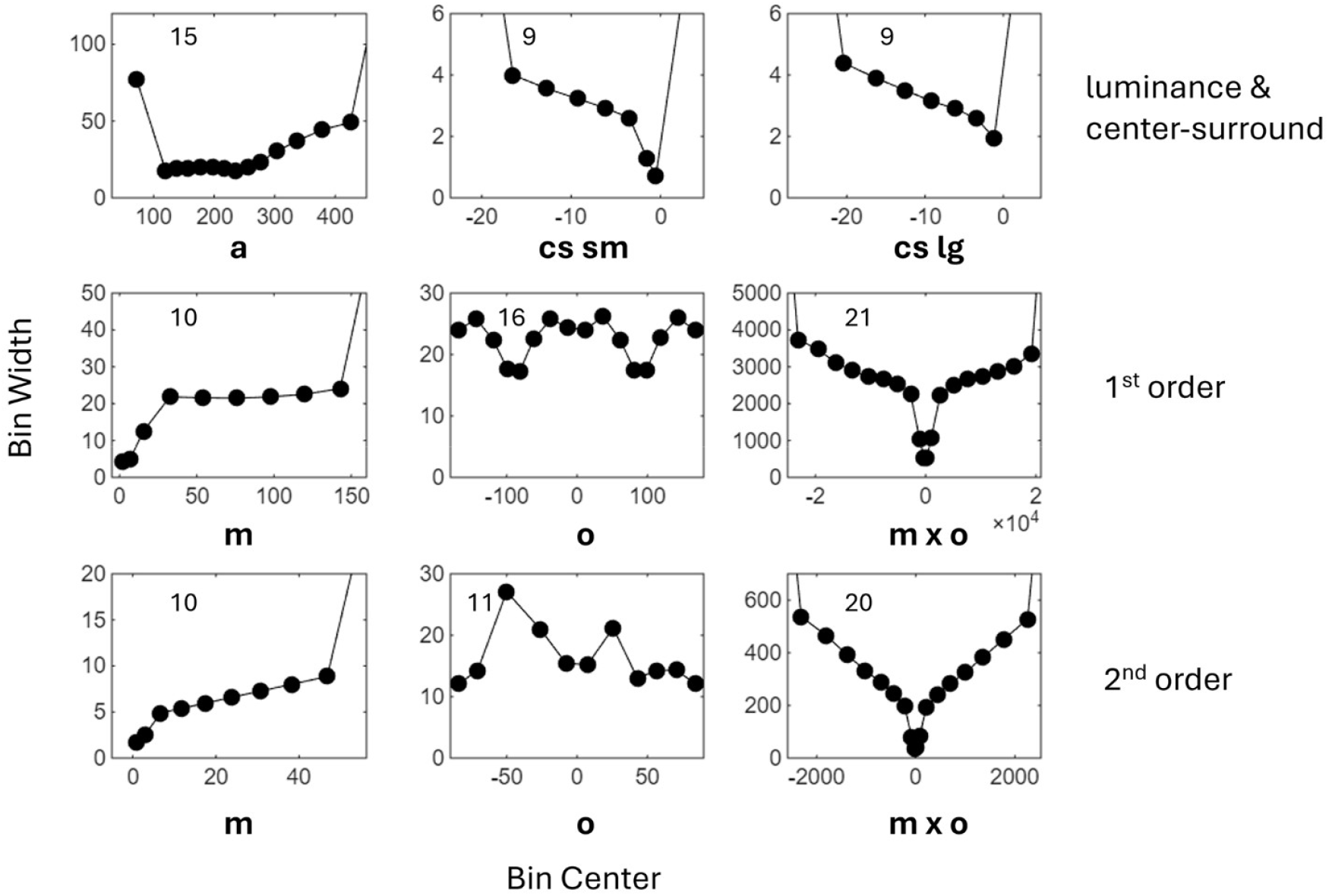
Optimal histogram bins estimated for nine feature dimensions using adaptive histogram equalization. Trained on Brodatz and Fabric textures. **Upper row**: achromatic color channel and center-surround filters. **Middle row**: derivative of Gaussians steerable filter (magnitude, orientation, product features). **Lower row**: second derivative of Gaussian steerable filter.

### Power log likelihood ratio

Let **P**_*a*_ and **P**_*b*_ be the vectors of observed power, at each possible spatial frequency**u**_*i*_ = (*u*_*i*_, *v*_*i*_), for patches *a* and *b*. Similar to above, the log likelihood ratio is given by

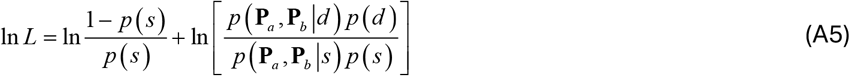

Assuming independent exponential distributions for the power at each frequency (which we verified for natural textures) we have

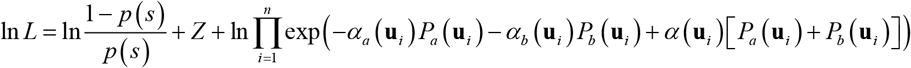

where,

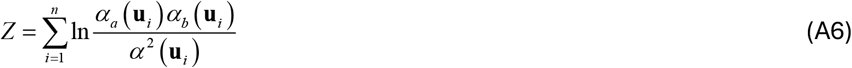

The maximum likelihood estimates of the exponential distribution parameters are given by

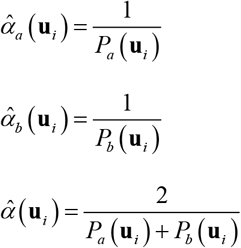

Thus,

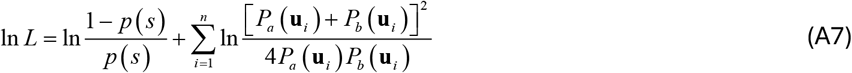

We found that performance was improved by including a weak-response (noise) suppression constant *β* :

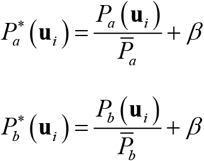

where 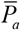 and 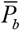 are average powers in the two patches.

With this addition, the log likelihood ratio becomes:

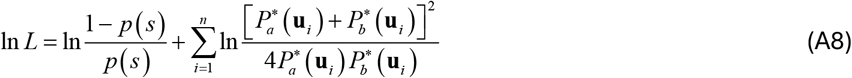

### Steerable filters

First and second derivative of Gaussian steerable filters (Freeman & Adelson,1991) were used to measure the edge and bar information. The advantage of steerable filters is that filtering a patch with just two or three kernels is sufficient to directly estimate, at every pixel location, the filter orientation that gives the maximum response as well as the value of that maximum response.

The first derivative steerable filter is given by

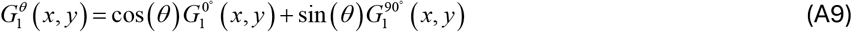

where *θ* is orientation, 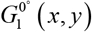 is the derivative of a Gaussian in the *x* direction and 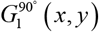 is the derivative of the Gaussian in the *y* direction. To compute the orientation of the max response and the max response at every pixel, one first convolves the image with the two kernels

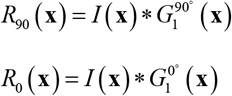

where **x** = (*x, y*). The estimated orientation and max response at each pixel are then given by the following two equations:

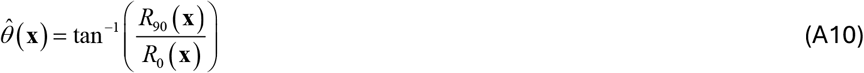

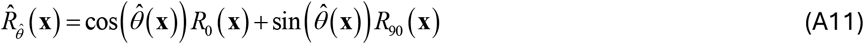

The second derivative steerable filter is given by

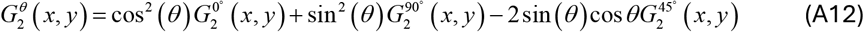

where the kernels are the second derivative of the Gaussian in the *x* direction, *y* direction, and *x = y* direction. As above, one first convolves the image with each of the three kernels. The formulas for directly estimating the orientation and max response at each pixel location from the three filtered images are a little trickier and may not be well known. Convolving both sides of Eq. A12 with the image gives

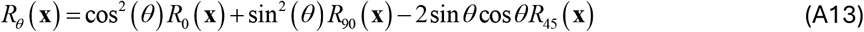

Taking the derivative with respect to *θ* and setting equal to zero gives

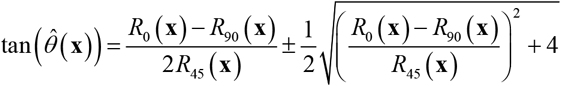

which gives two estimates of *θ* and two estimates of *R*_*θ*_ :

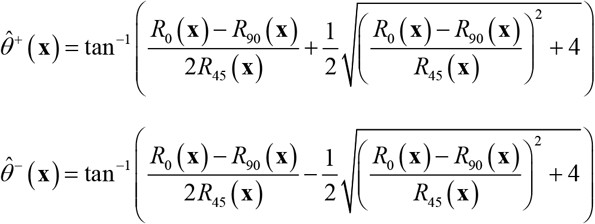

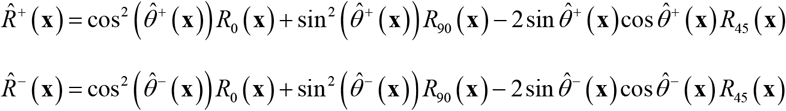

The estimated orientation is given by

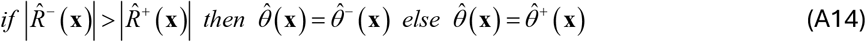

and the estimated response is

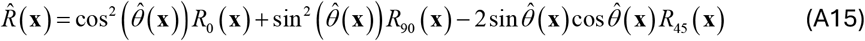

Because the texture patches are quite small, we kept the size of the steerable filters as small as possible: 3 × 3 pixels with a standard deviation of 1 pixel. Fig. A2D shows the spatial frequency tuning of the two steerable filters and the two center-surround filters. The peak spatial frequency in the fovea is 8 c/deg for the first derivative filter, 11 c/deg for the large center-surround filter and 21 c/deg for the second-derivative and small center-surround filters. These frequencies drop by a factor of two at each successive step in eccentricity tested in the experiments (e.g., peaks of 1, 1.4, and 2.6 c/deg at the 11.55° eccentricity).

**Figure A2.**
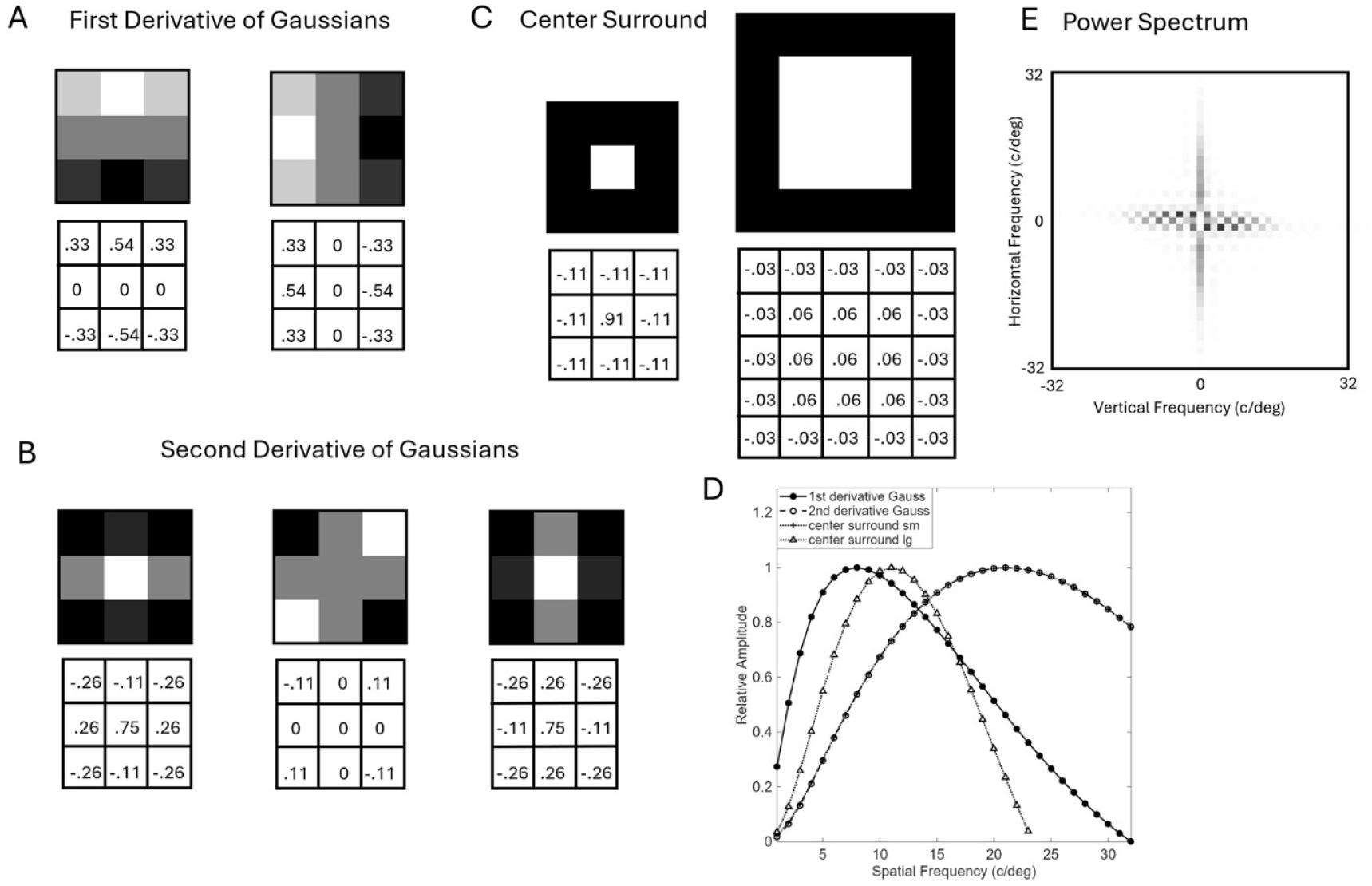
Task-independent features. **A**. Kernels of the first-derivative steerable Gaussian filter. Numbers sre the weights in each pixel **B**. Kernels of the second-derivative Gaussian steerable filter. **C**. Kernals of the center-surround filters. **D**. Spatial frequency tuning curves of the 4 filters in the fovea combined with the effect of the optical transfer function of the human eye. Despite the small kernal sizes of the steerable filters (3 × 3), there is only a small bias in the estimations of the magnitude and orientation of sinewaves. In other words, the steerablity is quite accurate. **E**. Patch power spectrum.

The small kernel sizes mean that the filters were not perfectly steerable. However, the bias in the estimation of magnitude and orientation across orientation of sinewave gratings was quite small. For the 1^st^ and 2^nd^ derivative filters the maximum bias in magnitude across orientation was about 5%. For the 1^st^ derivative filter the maximum bias in orientation was about 1.2° and for the 2^nd^ derivative filter about 1.7°. In other words, despite the small kernel size the filters were accurately steerable. To achieve this with the 2^nd^ derivative filter it was necessary to scale the 45° (middle) kernel to an energy of 0.22 rather than 1.0 like the other kernels.

### Task-dependent feature definitions (one-shot learning)

Fig. A3 lists all the log likelihood decision variables computed in the discrimination and segmentation tasks. The individual log likelihoods of same vs. different for the spot and contour features are given by a histogram decision variable (the first equation), with the bin bounds shown in Fig. A1. The power spectrum features decision variable is given by the fourth equation. All the other log likelihood decision variables follow from the optimal decision variables and bounds trained on Brodatz and Fabric textures using our software [https://github.com/abhranildas/Integrate-and-Classify-Normal-Distributions].

**Figure A3.**
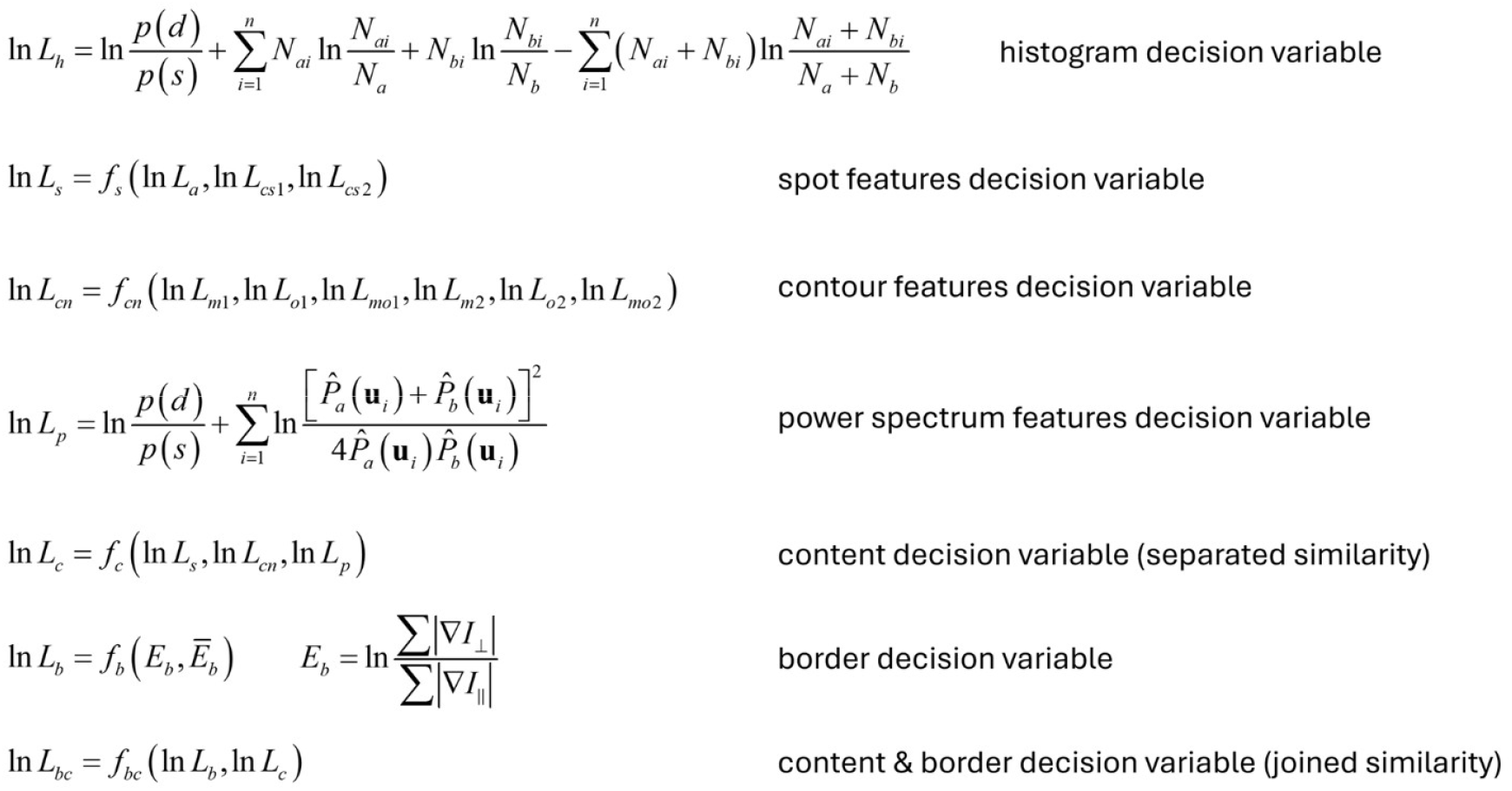
Task-dependent features computed on pairs of patches in the discrimination task and in the image segmentation task. The similarity measure for separated patches is based only on the content features of the patches. The similarity measure for joined patches is based on both the content and border features.

### Effect of retinal eccentricity

We modeled the effect of retinal eccentricity at the four eccentricities tested in the same-different texture discrimination experiments by down-sampling the input images, while keeping the filter kernel sizes (in pixels) fixed. The eccentricities correspond to approximate retinal eccentricities where the spacing of midget retinal ganglion cell receptive fields in humans drops by successive factors of two (e.g., the spacing at 12° eccentricity is approximately 1/8 of that in the fovea). The spacing values were estimated from the data of Curio & Allen (1990) for the horizontal meridian, using the algorithm of Watson (2014). For each factor-of-two reduction in sampling we first blurred with a 3 × 3 Gaussian kernel and then down-sampled by a factor of two using the nearest-neighbor method. This procedure keeps the loss of information due to down-sample accurate, while minimizing aliasing.

#### Texture sheets

**Figure A4.**
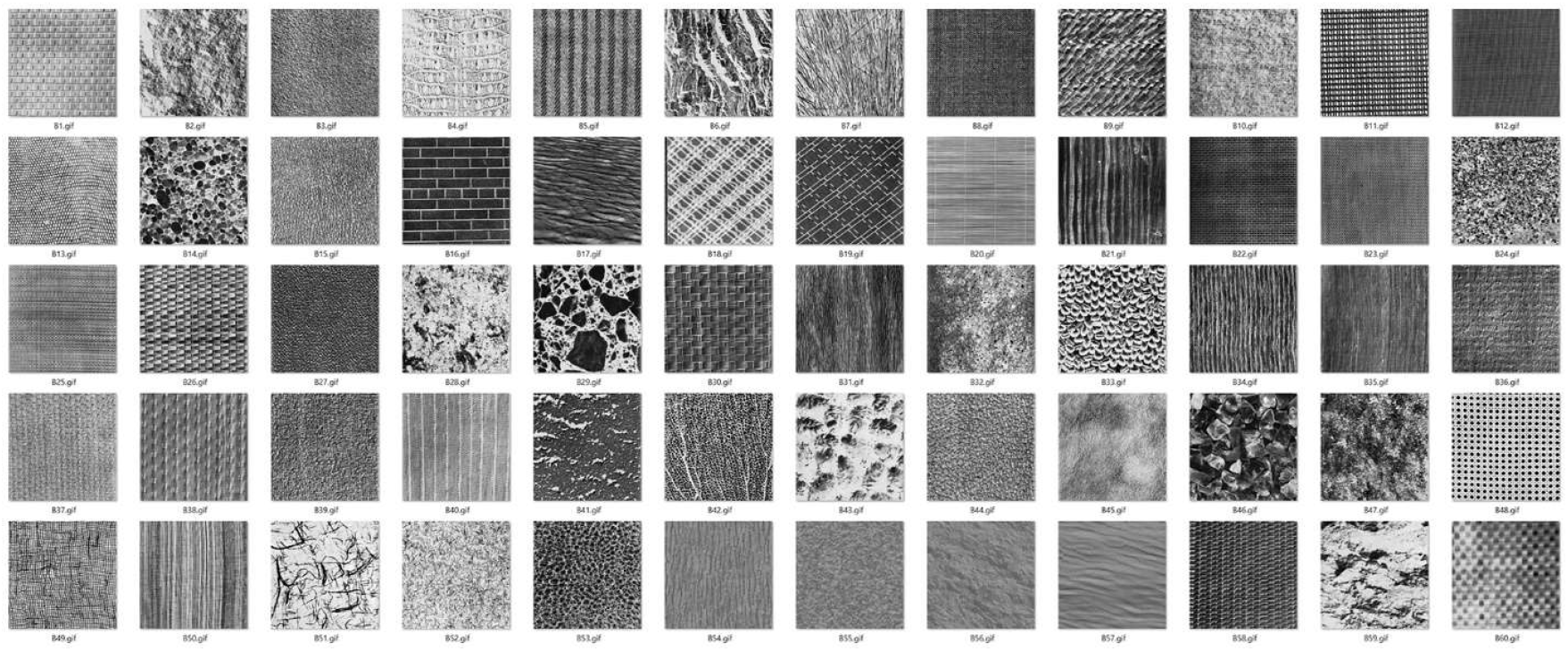
Thumbnails of the Brodatz texture sheets. Actual image size 640 × 640.

**Figure A5.**
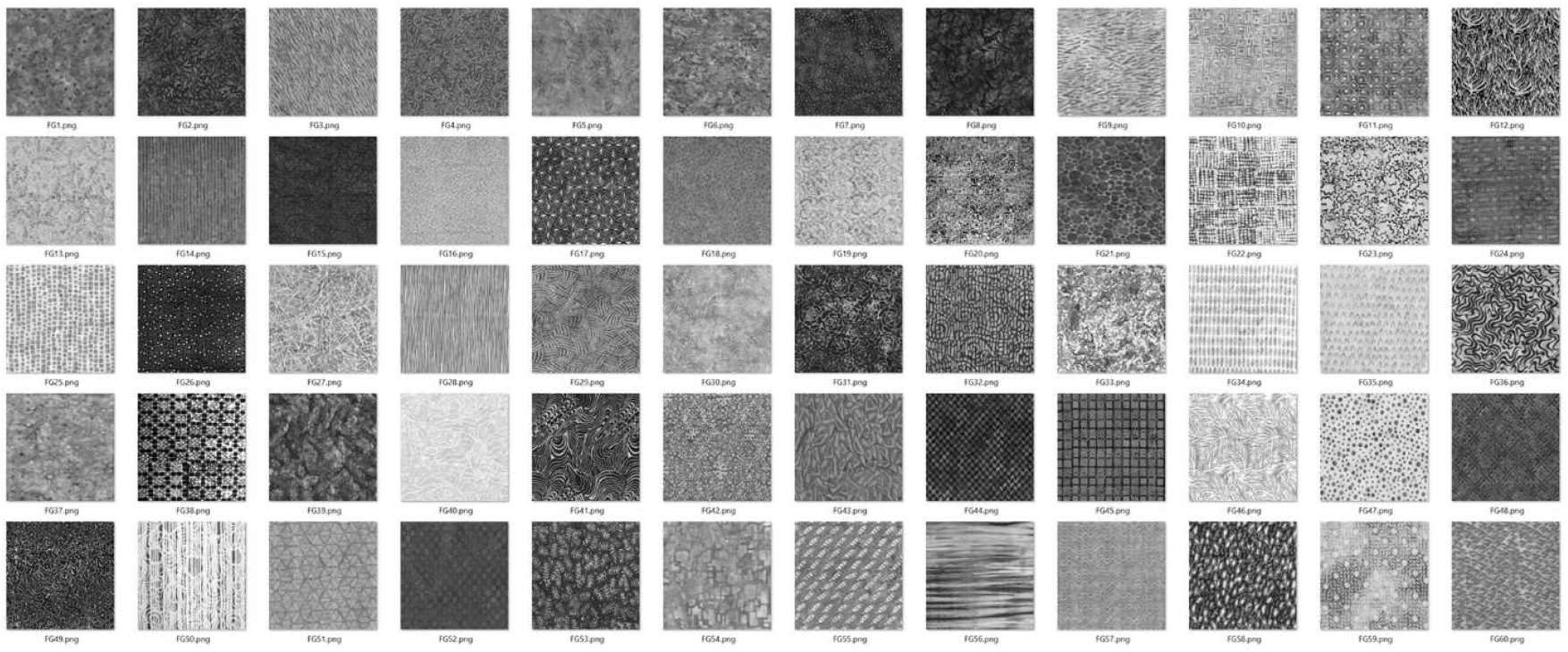
Thumbnails of the Fabric texture sheets. Actual image size 640 × 640.

